# Stochastic time-dependent enzyme kinetics: closed-form solution and transient bimodality

**DOI:** 10.1101/2020.06.08.140624

**Authors:** James Holehouse, Augustinas Sukys, Ramon Grima

## Abstract

We derive an approximate closed-form solution to the chemical master equation describing the Michaelis-Menten reaction mechanism of enzyme action. In particular, assuming that the probability of a complex dissociating into enzyme and substrate is significantly larger than the probability of a product formation event, we obtain expressions for the time-dependent marginal probability distributions of the number of substrate and enzyme molecules. For delta function initial conditions, we show that the substrate distribution is either unimodal at all times or else becomes bimodal at intermediate times. This transient bimodality, which has no deterministic counterpart, manifests when the initial number of substrate molecules is much larger than the total number of enzyme molecules and if the frequency of enzyme-substrate binding events is large enough. Furthermore, we show that our closed-form solution is different from the solution of the chemical master equation reduced by means of the widely used discrete stochastic Michaelis-Menten approximation, where the propensity for substrate decay has a hyperbolic dependence on the number of substrate molecules. The differences arise because the latter does not take into account enzyme number fluctuations while our approach includes them. We confirm by means of stochastic simulation of all the elementary reaction steps in the Michaelis-Menten mechanism that our closed-form solution is accurate over a larger region of parameter space than that obtained using the discrete stochastic Michaelis-Menten approximation.

## 1 Introduction

The mechanistic basis of the simplest single-enzyme, single-substrate reaction consists of a reversible step between an enzyme and a substrate, yielding the enzyme–substrate complex, which subsequently forms the product. This reaction is commonly called the Michaelis-Menten (MM) reaction [1, 2].

For over a century, the dynamics of this reaction have been extensively studied using deterministic rate equations. Because these equations do not admit an exact closed-form solution, various approximations have been devised to obtain insight into the underlying dynamics. Use of the quasi-equilibrium or quasi steady-state approximations lead to the famous Michaelis-Menten equation, an ordinary differential equation relating the rate of product formation and the substrate concentration (see [3] for a discussion of these approximations and their range of validity). This equation provides a simple means to extract the relevant kinetic parameters (the Michaelis-Menten constant and the maximum velocity) from experimental data. The Michaelis-Menten equation has also been solved exactly leading to explicit expressions for the time-evolution of the substrate (and product) concentration [4].

The stochastic formulation of enzyme kinetics, while not as much studied as its deterministic counterpart, has received increasing attention since the 1960s when the chemical master equation (CME) for the MM reaction mechanism was first derived and studied by Anthony F. Bartholomay [5]. The CME is a probabilistic discrete description of chemical reaction kinetics that is valid in well-mixed environments for point reacting particles [6,7]. Its relevance lies in its ability to describe kinetics when the molecule numbers are low, conditions typical in intracellular environments, e.g., the median copy number per cell of most enzymes in *E. coli* is below a thousand [8]. Research efforts in the area of stochastic chemical kinetics can be, broadly speaking, categorized into three types: (i) The search for a solution of the CME for the MM reaction and its various extensions, i.e., obtaining a closed-form solution for the time-dependent or steady-state probability distribution of the molecule numbers of each species in the reaction system [9, 10]. (ii) The reduction of the CME and the construction of the stochastic equivalent of deterministic approximations (such as the fast equilibrium, quasi steady-state and total quasi steady-state approximations) and understanding their regime of validity [11–24]. (iii) The derivation of exact or approximate expressions for the mean of the stochastic rate of product formation and an investigation of the differences or similarities from the predictions of the deterministic Michaelis-Menten equation [25–31].

The majority of the literature has focused on (ii) and (iii). There are very few studies that focus on (i) principally because the CME is notoriously difficult to solve analytically [22]. In this paper, we are interested in deriving new solutions of the CME for enzyme kinetic systems and hence next we briefly review the known solutions (see also [32] for a lengthier discussion). Arányi and Tóth [9] were the first to exactly solve the CME introduced by Bartholomay for the special case in which there is only one enzyme molecule with several substrate molecules in a closed compartment; in particular, they obtained an exact expression for the joint distribution of the number of substrate and enzyme molecules as a function of time (since the original paper is rather difficult to find, in Appendix A we have reproduced the derivation in a concise manner). Another exact solution is reported in [10] by Schnoerr et al. who derive the exact steady-state solution for the CME describing the MM reaction system with one enzyme molecule and augmented with a substrate production reaction step (to model for example the translation of substrate). To our knowledge, there are no known exact or approximate solutions for the time-dependent probability distribution solution of the CME of the MM reaction system with multiple enzyme molecules; however, expressions for the mean rate of product formation have been derived and some of these papers are referenced in point (ii) above.

In this paper, our aim is to (a) derive an expression for the approximate time-dependent solution of the CME of the MM reaction system with multiple enzyme molecules under quasi-equilibrium conditions; (b) compare and contrast this solution with the solution of an often used reduced CME for the MM reaction in the literature; (c) use the closed-form solution to identify interesting dynamical phenomena. Our paper is divided as follows. In Section 2, we briefly review the main results known for deterministic enzyme kinetics, focusing in particular on the quasi-equilibrium approximation. In Sections 3.1 and 3.2, we introduce our method by first applying it to the MM reaction with a single enzyme molecule and subsequently to the case of multiple enzyme molecules. The method consists of three steps: (1) using a time scale separation method called averaging [33] to define groups of rapidly equilibrating states which then allows the derivation of a master equation describing the Markovian dynamics of these groups on the slower time scale; (2) solving the resultant time-dependent, single variable master equation for the group dynamics using the method developed in [34] which has the advantage of bypassing the calculation of the eigenvectors of the transition matrix and hence considerably simplifies the analytical computations; (3) using the time-dependent solution describing the group dynamics to construct the marginal time-dependent distributions for both the numbers of substrate and enzyme molecules. We use the closed-form solution to find the regions of parameter space where transient bimodality of the distribution of substrate molecules occur. In Section 4, we show that our solution is accurate over a wider region of parameter space than the solution of a commonly used reduced master equation with a propensity that has the same hyperbolic dependence on the number of substrate molecules as the deterministic Michaelis-Menten equation (an approach popularized by Rao and Arkin [12]). In Section 5, we show that the same three-step method used in Sections 3.1 and 3.2, can be used to derive time-dependent distributions for multi-substrate enzyme reactions. We finish by discussing our results in Section 6.

## 2 Deterministic enzyme kinetics

Before progressing to stochastic enzyme kinetics we first briefly outline some of the main results known for deterministic enzyme kinetics. We consider the chemical reaction system:

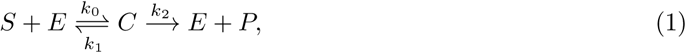

where *S* denotes the substrate species, *E* denotes the enzyme species, *C* denotes the enzyme-substrate complex and *P* denotes the product. This system can be thought of as a reduction of the more biologically realistic set of reactions:

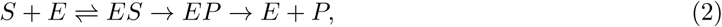

where the unbinding of the product from the enzyme is very fast. Without loss of generality, we assume the initial condition for this system is that all enzymes are unbound to the substrate. There are two conservation laws for this system: [*E*] + [*C*] = [*E*]_0_ and [*S*] + [*C*] + [*P*] = *N*, where [*i*] denotes the concentration of species *i*, [*i*]_0_ denotes the initial concentration of species *i* and *N* is the initial substrate concentration. Assuming well-mixed conditions and the law of mass action, the deterministic dynamics of the reaction system in Eq. (1) are described by a set of coupled ordinary differential equations (commonly called the rate equations) describing the time-evolution of the substrate and complex concentrations:

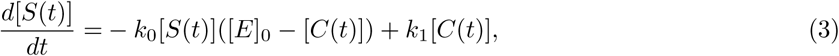

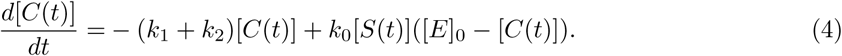

Note that the time-dependent concentrations of *E* and *P* can be straightforwardly obtained from the time-dependent solutions of *C* and *S* by means of the conservation laws previously stated. Although seemingly simple, Eqs. (3) and (4) are not easy to solve analytically for the time-dependent analytic solution, and as such one is limited to finding approximate solutions. Two of the most common approximations used in the literature are the (i) *quasi steady-state assumption* (QSSA) and (ii) the *quasi-equilibrium approximation* (QEA), also called the rapid equilibrium approximation or the reverse quasi steady-state assumption. The QSSA, derived by Briggs and Haldane [35], assumes that after a short transient, the concentration of the complex (and enzyme) is in a quasi steady-state (with regard to the substrate and product); thus under the QSSA, it is assumed that *d*[*C*(*t*)]*/dt* ≈ 0. See [36] for a detailed discussion of this approximation and for its range of validity. On the other hand, the QEA assumes that substrate binding and dissociation occur much more rapidly than product formation such that the substrate, enzyme and complex are approximately in equilibrium. Thus under the QEA, it is assumed that *d*[*S*(*t*)]*/dt* ≈0; this approximation, popularized by Michaelis and Menten [1], is commonly used in the analysis of various biochemical models [37].

Enforcing either the QSSA or QEA leads to the following effective rate equation describing the time-evolution of the substrate concentration:

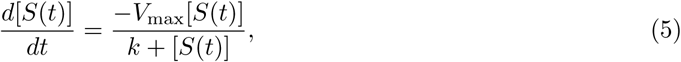

where *V*_max_ = *k*_2_[*E*]_0_, *k* = (*k*_1_ + *k*_2_)*/k*_0_ if the QSSA is used, *k* = *k*_1_*/k*_0_ if the QEA is used, and where the conservation law [*S*] + [*P*] = *N* holds. Eq. (5) has been solved perturbatively in a number of studies, all of which also assessed the validity of the QSSA [36, 38]. An exact solution was reported in [4] which is given by:

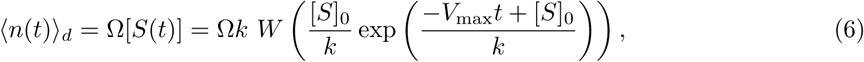

where ⟨*n*(*t*) ⟩_*d*_ gives the (deterministic) mean *number* of substrate molecules at time *t*, Ω is the volume of the system, and *W* (·) is the principal branch of the Lambert *W* function (also known as the Omega function). In the rest of this article, we study the stochastic equivalent of the QEA and thus we shall use *k* = *k*_1_*/k*_0_.

## 3 Stochastic QEA analysis

### 3.1 Single enzyme

For simplicity, we first illustrate the method by solving the enzyme system described in Eq. (1) for the case of one enzyme molecule with initially *N* substrate molecules. Since there are no birth-death processes coupled to any species, the conservation equations *n*_*E*_ + *n*_*C*_ = 1 and *n* + *n*_*C*_ + *n*_*P*_ = *N* hold, where *n* denotes the number of substrate molecules and all other *n*_*i*_ denote the number of species *i*. Without loss of generality, we set the size of the system to Ω = 1 for the rest of the paper.

We label the microstate of the reaction network in Eq. (1) as (*n, n*_*E*_), which fully specifies the state of the system due to the conservation laws stated previously. The possible transitions between all of the discrete microstates of this system are illustrated in Fig. 1(i): the system starts from the state (*N*, 1) and eventually ends up in the state (0, 1). Our goal now will be to find the marginal probability distribution *P* (*n*; *t*), i.e., the probability of observing *n* substrate molecules at a time *t*.

**Figure 1:**
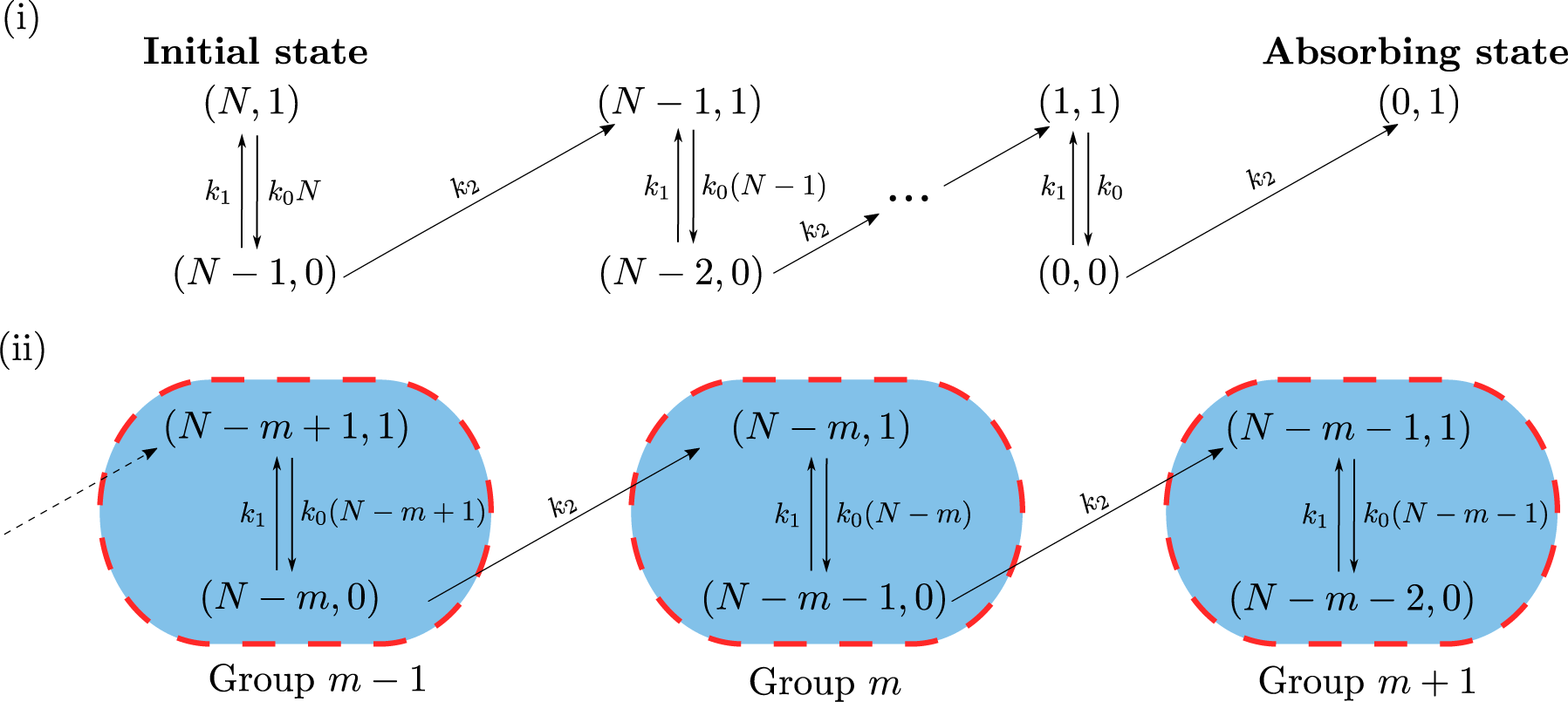
Illustration of the enzymatic system described by a single enzyme and *N* initial substrate molecules. (i) Markovian dynamics of the enzyme kinetic system described by a single enzyme. The initial condition for the system is (*N*, 1), and as *t* → ∞ the microstate of the system is guaranteed to be that of the absorbing state (0, 1), with no remaining substrate and one free enzyme. (ii) Markovian dynamics in the reduced model, where processes occurring in a group are assumed to be much faster than the interactions between the groups themselves. The label ‘group *m*’ denotes the set of fast processes that have *N* − *m* substrate molecules when the enzyme is free; hence, it is easily seen that there are *N* + 1 groups in total with labels *m* = {0, 1, 2, …, *N* − 1, *N*}.

Assuming Markovian dynamics [22], it follows that the time-evolution of *P* (*n, n*_*E*_; *t*) (the probability of observing *n* substrate molecules and *n*_*E*_ enzyme molecules at a time *t*) is given by the CME:

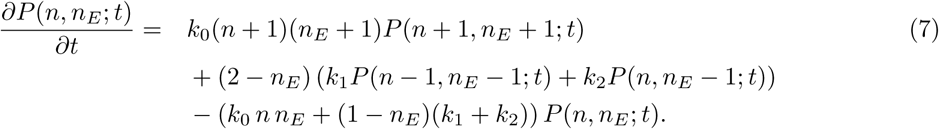

The standard approach involves introducing the time-dependent marginal generating functions 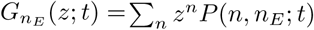 and attempting to solve the generating function partial differential equations, e.g., using eigenfunction methods [7]. However, this standard method quickly leads one to mathematical difficulty. An analytic solution only presents itself in a non-cumbersome form where one assumes the initial state contains a single substrate molecule [9]. In Appendix A we summarise the single enzyme solution provided by [9], and its complexity even in the single substrate molecule case motivates the analysis we present below.

We take a different approach. We first simplify the problem through the use of averaging [33,39,40]. Specifically the procedure lumps together microstates equilibrating on a fast timescale in groups which then allows one to write a master equation describing the dynamics of the groups on the slow timescale. We shall assume that the slow timescale is that associated with product formation, i.e. *k*_2_ is sufficiently small (we will be more precise what this really means later) and hence the averaging procedure is in the same spirit as the QEA discussed in Section 2.

Since *k*_2_ is small, it follows that we can group all microstates that are in rapid equilibrium with each other (due to the fast processes of binding and unbinding of substrate from the enzyme) as shown in Fig. 1(ii); group *m* is then the set of fast processes that contain *N* − *m* substrate molecules when *n*_*E*_ = 1 and *N* − (*m* + 1) substrate molecules when *n*_*E*_ = 0. We define 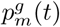 as the probability to be in group *m* at a time *t*, and 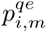 as the probability that if the system is in group *m* it has *i* free enzymes. Once these probabilities are found, we can construct *P* (*n*; *t*), based on the fact that there are two microstates that contain *n* substrate molecules: (*n*, 0) and (*n*, 1) associated with groups *N* − (*n* + 1) and *N* − *n* respectively. This means that:

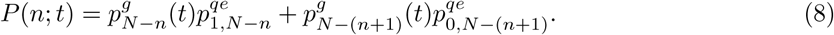

In the case of the single enzyme system studied in this section, the quasi-equilibrium probabilities are trivial (since there are only two microstates in each group) and are given by:

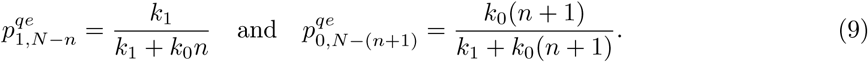

All that remains is the task of finding 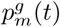. To do this we first write the master equation for the transitions between groups. Rescaling time as *t*′ = *k*_2_*t* and making use of the previous definition, *k* = *k*_1_*/k*_0_, the master equation for the groups is:

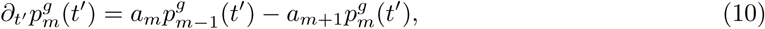

where:

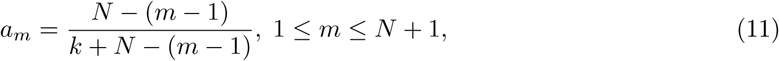

and *a*_*i*≤0_ = 0. Note that *a*_*m*_ is the probability of the jump from group *m* −1 to group *m* in a unit interval of rescaled time. From Fig. 1 the probability of the jump from group *m* −1 to group *m* in a unit interval of normal time is equal to *k*_2_ multiplied by the probability of being in the microstate (*N* −*m*, 0) which under the rapid equilibrium assumption is *k*_0_(*N* −*m* + 1)*/*(*k*_1_ + *k*_0_(*N*− (*m* −1))). Due to time rescaling, the factor of *k*_2_ disappears and hence follows Eq. (11).

Since there are *N* + 1 groups in total, Eq. (10) corresponds to a system of *N* + 1 ODEs which can be concisely written as the matrix equation:

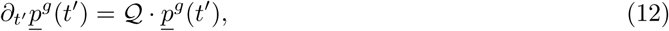

where *<u>p</u>*^*g*^(*t*′) is a *N* + 1 element column vector defined by 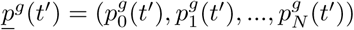 and 𝒬 is a (*N* + 1) *×* (*N* + 1) lower-bidiagonal square matrix defined by:

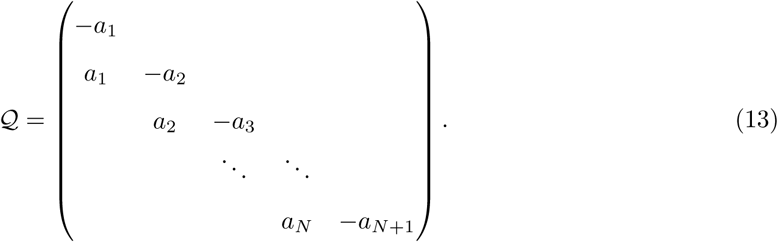

As we will describe below, we solve the set of ODEs given by Eq. (12) using the method described in [34] which provides an *exact* time-dependent solution for any one-variable one-step master equation with finite number of microstates *as long as one can find the eigenvalues of the transition rate matrix exactly*. In our case, the eigenvalues of 𝒬 are trivial, since 𝒬 is lower-bidiagonal, and they are given by the diagonal elements. Hence the eigenvalues of 𝒬 are given by *λ*_*i*_ = −*a*_*i*_, 1 ≤ *i* ≤ *N* + 1. Note that *λ*_*N*+1_ = 0 and is the largest eigenvalue, with all *λ*_1≤*i*≤*N*_ < 0 as is required by the Perron-Frobenius theorem for Markovian systems that are ergodic [41].

We now proceed to use these eigenvalues to find the time-dependent solution to Eq. (12). The solution to this set of ODEs is formally given by:

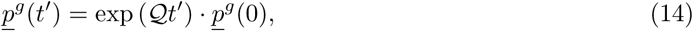

where exp(𝒬*t*′) is defined as a matrix exponential. For a general master equation, this matrix exponential is typically difficult to deal with, however in our case 𝒬 is lower-bidiagonal and hence we can proceed via the method of [34]. We first consider Cauchy’s integral formula for matrices, explicitly given by [42]:

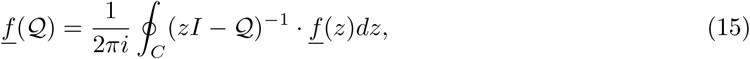

where *C* is a closed contour in the complex plane that encloses all the eigenvalues of 𝒬 and *I* is the identity matrix. Taking 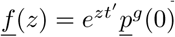 we then arrive at:

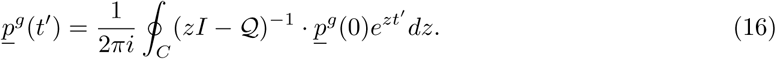

A typical initial condition is 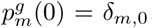, meaning that we always start in group 0 which contains the microstates (*N*, 1) and (*N* – 1, 0), as is shown in Fig. 1(i). Note that *δ*_*i,j*_ is the Kronecker delta. Using this initial condition, Eq. (16) becomes:.

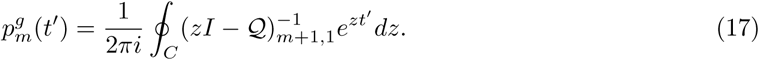

We show at the end of this section how to extend the time-dependent solution for a general initial distribution. Since it is bidiagonal, the inverse of *zI* − 𝒬 can easily be found via Cramer’s rule [43]:

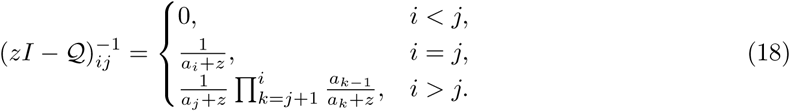

Substituting this into Eq. (17) then gives us:

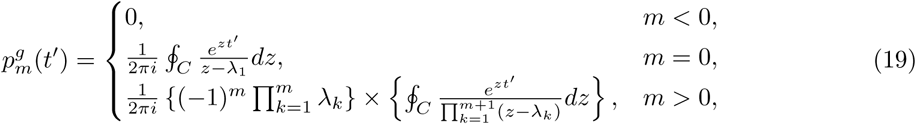

where we have utilised the relation *λ*_*i*_ = −*a*_*i*_. These integrals can then be evaluated using Cauchy’s residue theorem [44], explicitly stated as:

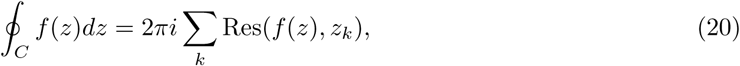

where the values *z* = *z*_*k*_ are poles of *f* (*z*) within *C* and the residues are Res(*f* (*z*), *z*_*k*_) = lim_*z*→*zk*_ (*z* − *z*_*k*_)*f* (*z*) for the simple poles in Eq. (19). Note that the poles of the complex integrals in Eq. (19) are the eigenvalues of 𝒬. Therefore, from Eq. (19) we finally get an expression for 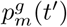 as:

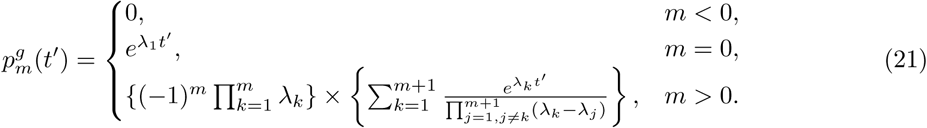

Hence the time-dependent probability distribution *P*(*n*; *t*) is given by Eq. (8) together with Eqs. (9) and (21). The extension to a more general initial distribution is then relatively simple. Consider some initial distribution *<u>p</u>*^*g*^(0) = *<u>q</u>*, where *<u>q</u>* is an *N* +1 element vector; the time-dependent group probability 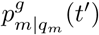 is then given by the weighted sum:

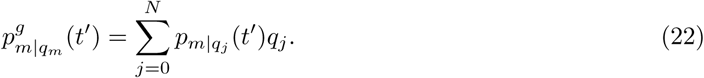

This initial condition could be useful to model variation in the initial number of substrate molecules due to uncertainty introduced by experimental error or else due to the intrinsic noise in the reaction mechanism generating the substrate. Note that if *q*_*m*_ = *δ*_*m*,0_, one clearly recovers the analysis shown above. For the rest of the paper we only consider the initial condition 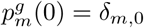, specifically where all enzymes are initially unbound to the substrate, but note that the analysis that follows can be easily extended for more general initial distributions.

In the beginning of this derivation, we stated that the main assumption is that *k*_2_ is sufficiently small. This statement can be made more precise as follows. From Fig. 1(ii) it is clear that the exit from group *m* can only occur when the enzyme is bound to substrate, i.e., from state (*N* − *m* − 1, 0). Now given that we are in this state, it follows that only two reactions can occur: either a reaction which causes a group change, i.e., (*N* −*m* −1, 0) → (*N* −*m* −1, 1) which occurs with rate *k*_2_ or a reaction that leads to no group change, i.e., (*N* − *m* − 1, 0) → (*N* −*m*, 1) which occurs with rate *k*_1_. Hence the probability of leaving the group is *k*_2_*/*(*k*_1_ + *k*_2_), from which follows that the microstates in each group will achieve quasi-equilibrium if *k*_2_ ≪ *k*_1_. Therefore, this is the condition under which our method provides a good approximation to the distribution of substrate molecules at all times.

We test the distributions predicted by Eq. (8) against the SSA in Fig. 2A(i-iii) and Fig. 2B(i-iii). In Fig. 2A(i-iii) we show that the solution is accurate for small *N* = 8, over a time range from *t*′ = 1 near the initial condition, to *t*′ = 12 close to the absorbing state, where the validity criterion *k*_1_ ≫ *k*_2_ holds. In Fig. 2B(i-iii) we observe that our solution agrees similarly well to the SSA for larger values of *N*. For a more general comparison of the exact solution to SSA through time, we can compute the mean and standard deviation from Eq. (8):

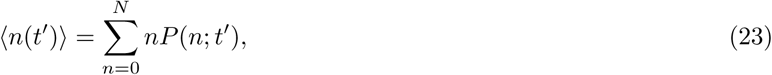

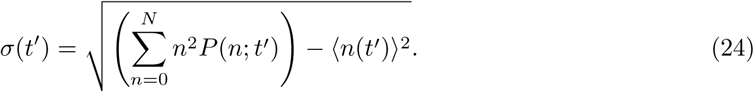

**Figure 2:**
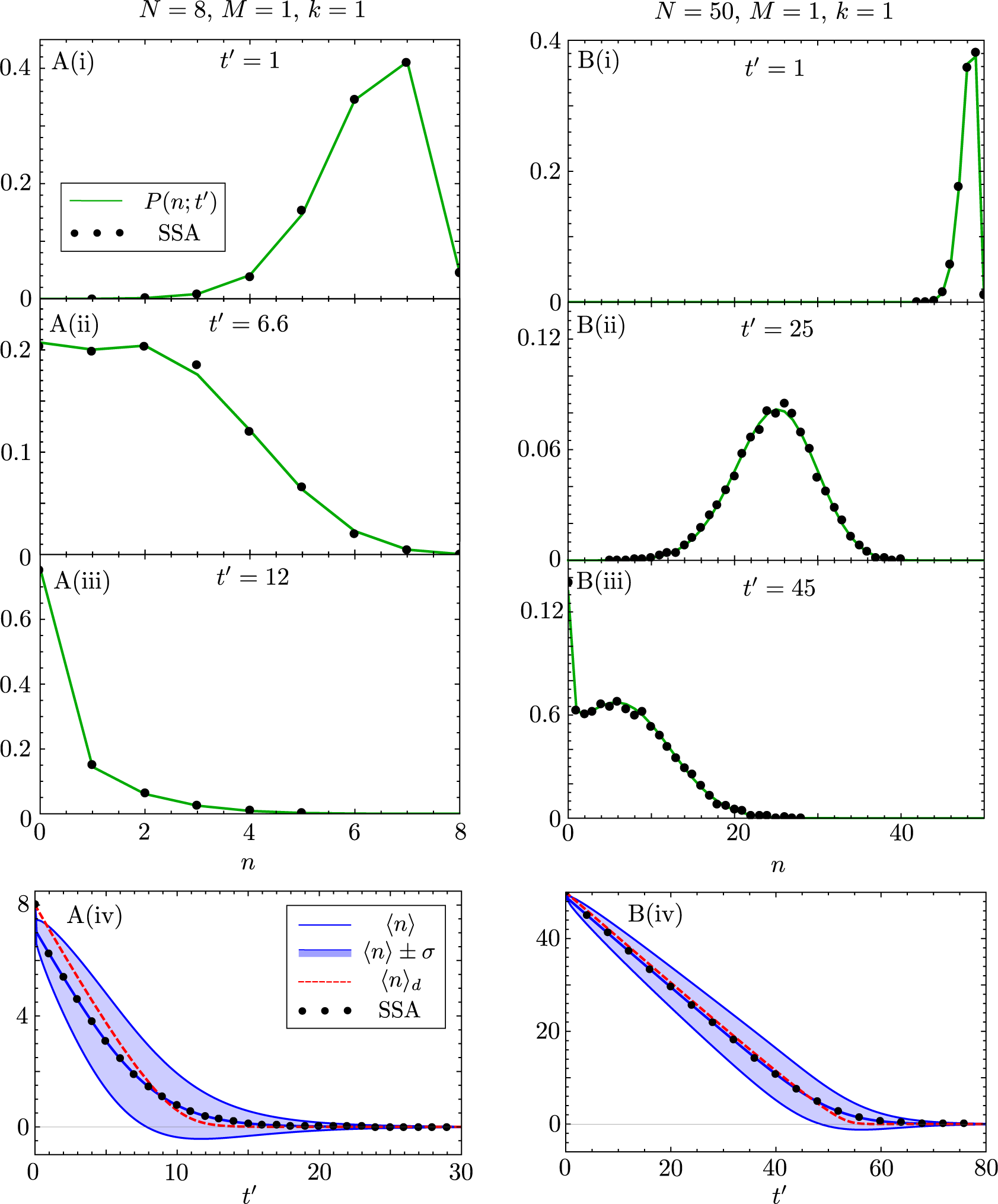
Comparison of the analytic time-dependent probability distribution of substrate molecules for the enzyme reaction in (1) with one enzyme molecule, i.e., *M* = 1, and *N* initial substrate molecules to the distribution obtained from the stochastic simulation algorithm (SSA) [6]. Note that the analytic solution is given by Eq. (8) together with Eqs. (9) and (21). In all cases we enforce *k*_1_*/k*_2_ ≫1 such that the quasi-equilibrium assumption behind the QEA is justified. We show the time-evolution of the distribution for substrate numbers, from near the initial condition to near the absorbing state, in two cases: A(i-iii) is for *N* = 8, *k*_0_ = *k*_1_ = 10^3^, *k*_2_ = 1, meaning that *k* = *k*_1_*/k*_0_ = 1. B(i-iii) is for *N* = 50, and all rate parameters as in the previous case. Note that the analytical solution (green lines) matches the SSA (black dots) for all times, for both a small and large initial number of substrate molecules. In A(iv) and B(iv) we show the corresponding plots of the time-evolution of the mean ⟨*n*⟩ and of the standard deviation σ of the distributions of substrate molecules, as predicted by our theory; these are compared with the mean calculated from the SSA and from the deterministic rate equation ⟨*n*⟩_*d*_ given by Eq. (6). Note that the deterministic mean is a better approximation to the stochastic mean for larger *N*. As shown in B(iii), and mildly in A(ii), the distribution can bimodal at intermediate times. Each SSA probability distribution is constructed from 10^5^ individual reaction trajectories.

The stochastic mean number of substrate ⟨*n*⟩ can then be compared to the deterministic solution mean number ⟨*n*⟩_*d*_ from Eq. (6).

In Fig. 2A(iv) we plot the evolution of the stochastic and deterministic mean substrate numbers in time, and compare them to the SSA for *N* = 8 and *k* = 1. We also show the standard deviation about the mean, i.e., ⟨*n*⟩ ±σ, where we have dropped the time dependence for brevity, given in the blue envelope. Clearly, ⟨*n*⟩ from Eq. (23) matches the mean predicted by the SSA for all times, whereas the deterministic mean *n* _*d*_ performs especially poorly (i) very soon after *t*′ = 0 and (ii) in the region where ⟨*n*⟩ _*d*_ ≦ 1. Both of these discrepancies are explained by the fact that deterministic analyses consider molecule number to be continuous. The explanation of (i) follows by considering the system after quasi-equilibrium has been reached between the states (*N*, 1) and (*N* − 1, 0) after a time 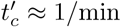{*k*_0_*N, k*_1_} which is small under the rapid equilibrium assumption. Because of the discreteness of the substrate molecules, ⟨*n*⟩ after a time 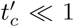 becomes an average over *n* = *N* and *n* = *N* − 1 weighted by the quasi steady-state probabilities 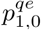 and 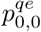 respectively, hence the step-like drop in ⟨*n*⟩ at 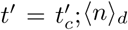 does not show this step-like drop at 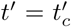 because it does not consider molecular discreteness, *hence the deterministic QEA does not capture the initial transient*. Averaging accounts for this initial transient, but not under the same definition as discussed in [4]; the initial transient captured here being the time between *t*′ = 0 and where quasi steady-state is achieved in group 0 at 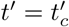 (this definition also follows for the next section on multiple enzymes). The explanation of (ii) follows since molecular discreteness is very important where ⟨*n*⟩ = 𝒪 (1), and properly accounting for it leads to differing dynamics for *n* in this region, whereas the behaviour of *n* _*d*_ does not change compared to ⟨*n*⟩_*d*_ ⪆ 1. As we shall see later, increasing the number of enzyme molecules removes this discrepancy between the stochastic and deterministic means, highlighting that the discrepancy seen here is because we do not consider enzyme molecules to be discrete in the deterministic analysis. Comparing Fig. 2A(iv) (with *N* = 8) and Fig. 2B(iv) (with *N* = 50), it is clear that as *N* becomes large, the deterministic mean becomes a better approximation of the true mean.

From Fig. 2B(iii) we observe that the distribution of substrate molecule numbers can be bimodal at intermediate times (there are two peaks at *n* = 0 and *n* = 6 at *t*′ = 45). This bimodality, though less conspicuous, can in fact be also observed in Fig. 2A(ii) with peaks at *n* = 0 and *n* = 2. From Fig. 2A(iv) and B(iv), we can see that in both cases the bimodality occurs at a time *t*′ when ⟨*n*⟩ − σ ≈ 0, i.e., when the fluctuations are large enough to cause frequent transitions to the absorbing state. This type of dynamical phase transition (which we shall refer to as transient bimodality), from a unimodal distribution to a bimodal one and then back to a unimodal one, as time progresses, has also been recently observed in genetic feedback loops [39] and is known in non-biological systems [45, 46]. We will discuss this phenomenon more extensively in later sections.

### 3.2 Multiple enzymes

We now extend the solution to the enzyme system (1) to the case where initially there are *N* free substrate molecules and *M* free enzyme molecules with the constraint of substrate abundance, i.e., *N* ≥*M*. Note that the solution to the system with *M* ≥ *N* follows as a special case of the *N* ≥ *M* system, discussed at the end of this section.

We proceed in solving this system as we did in the single enzyme case: assuming *k*_2_ is sufficiently small, we group the fast processes together to form *N* + 1 groups between which the transitions are significantly slower than those between the fast internal states of an individual group. The Markov chain describing the system split into groups is shown in Fig. 3. Our task is then to find (i) the equilibrium probabilities 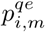 of being in each fast internal state *i* given we are in group *m* and (ii) to find the time-dependent probability 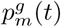 of being in group *m*. Knowledge of both (i) and (ii) will allow us to construct the distribution of interest, *P* (*n*; *t*).

**Figure 3:**
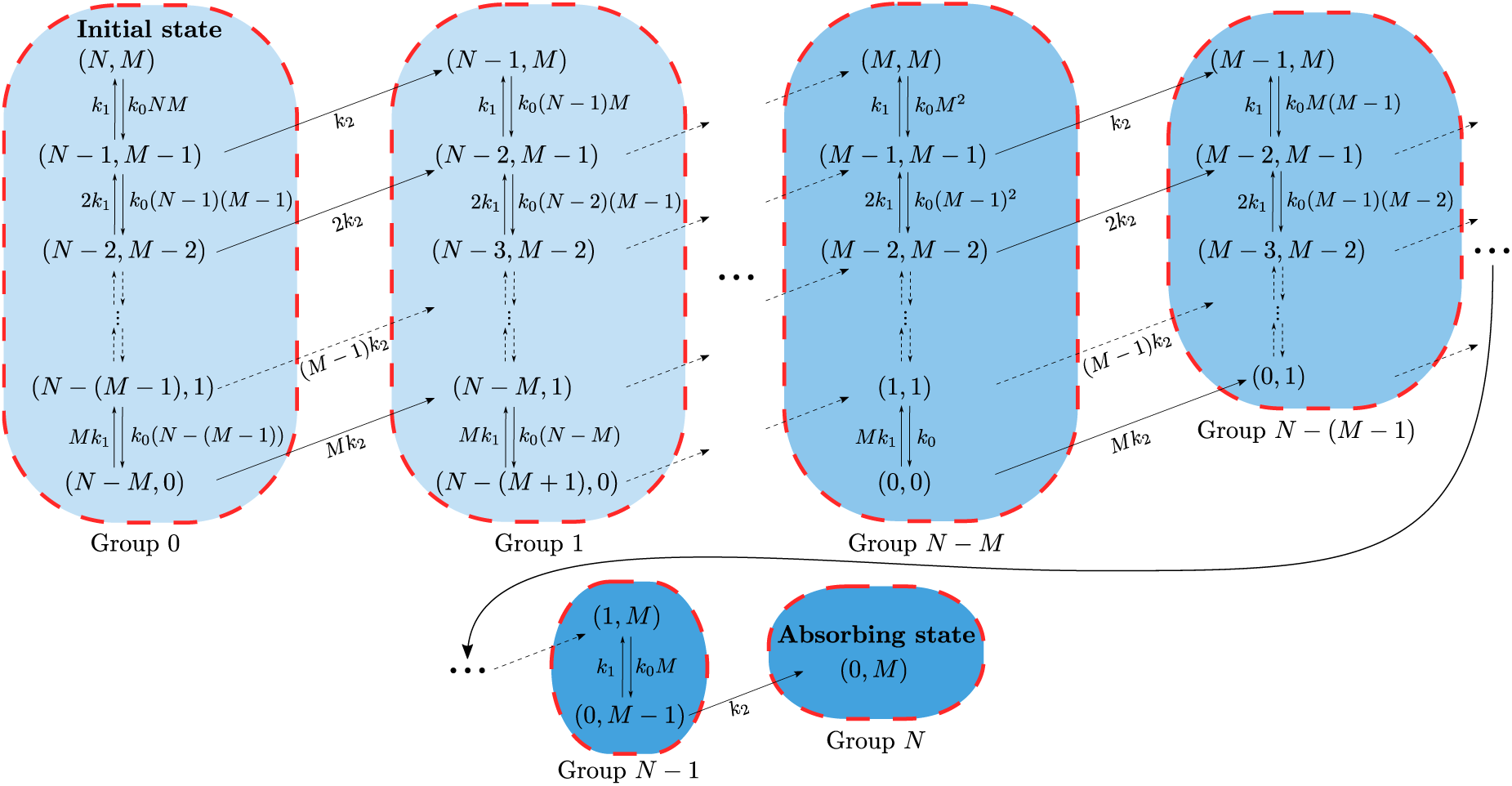
Illustration showing the transitions between the discrete microstates of the enzyme system (1) with initially *M* enzymes and *N* substrate molecules where *N* ≥ *M*. Fast processes are aggregated together, with each set of fast processes corresponding to a *group*. The label ‘group *m*’ denotes the set of fast processes with *N* − *m* free substrate molecules when *all* enzymes are free. Groups 0 ≤ *m* ≤ *N* − *M* have *M* + 1 fast internal states, whereas groups *N* − *M* < *m* ≤ *N* have *N* − *m* + 1 fast internal states. Note that as *t* → ∞ we are guaranteed to be in the absorbing state (0, *M*).

We begin by finding the probabilities 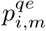 and revise its definition for the multiple enzyme case: 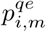 is the probability that if the system is in group *m* it has *M* − *i* free enzymes. Now, finding 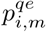 for any group 0 ≤*m* < (*N* − 1) is more complicated than was the case for a single enzyme system, since there we had only two fast internal states in each group. To proceed we consider the following Markovian dynamics of a system with *L* + 1 possible microstates:

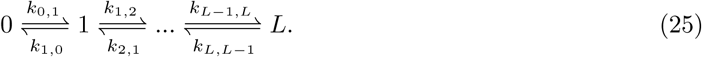

One can then write the master equation for this dynamical system in matrix form:

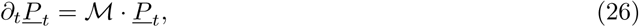

where *<u>P</u>*_*t*_ = (*P*_*t*_(0), *P*_*t*_(1), …, *P*_*t*_(*L*)), *P*_*t*_(*i*) is the probability of being in microstate *i* at time *t* and

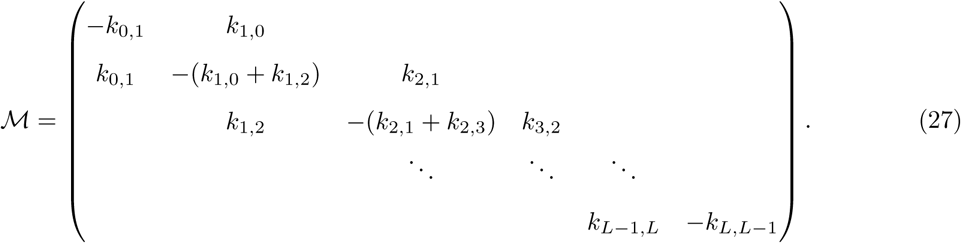

Enforcing the quasi-equilibrium condition, *∂*_*t*_(·) = 0, converts the system of *L* + 1 ODEs in Eq. (26) into a system of *L*+1 simultaneous equations in the equilibrium microstate probabilities *P* (*i*), given by *ℳ· <u>P</u>* = 0. One can explicitly solve this set of simultaneous equations under the constraint Σ_i_ *P* (*i*) = 1, yielding the probabilities:

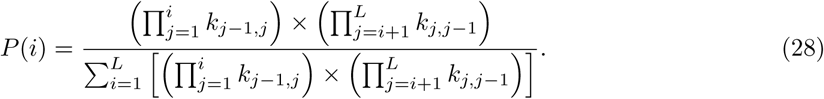

Using this result we can find the quasi-equilibrium probabilities for each group shown in Fig. 3. First, we consider the groups 0 ≤*m* ≤ *N* −*M*, each with *M* + 1 fast internal states as these groups contain more (or the same number) free substrate molecules than enzymes. Taking the specific example of group *m* = 0, we see that we have a total of *M* +1 microstates, i.e., *L* = *M, k*_*j*−1,*j*_ = *k*_0_(*N* −(*j*−1))(*M* −(*j*−1)) and *k*_*j,j*−1_ = *jk*_1_, with 1 ≤ *j* ≤ *M*. Identifying 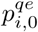 with *P* (*i*) in Eq. (28), we find that:

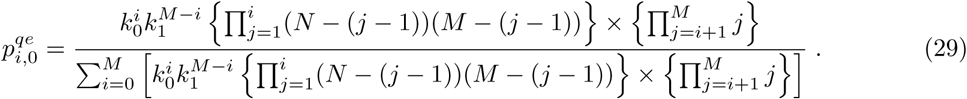

The result can be easily generalised for groups 0 ≤ *m* ≤ *N* − *M* and 0 ≤ *i* ≤ *M* :

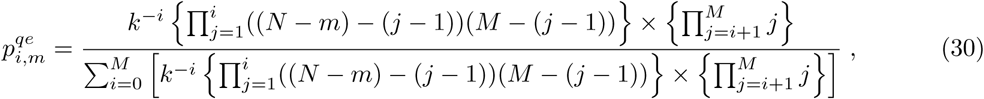

where we have re-introduced *k* = *k*_1_*/k*_0_. The dynamics of groups *N* −*M* < *m* ≤ *N* are slightly different as they contain fewer substrate molecules than enzymes. These groups correspondingly have *N* −*m* + 1 fast internal states, i.e., 0 ≤*i* ≤*N* −*m*. This leads to quasi-equilibrium probabilities of the form:

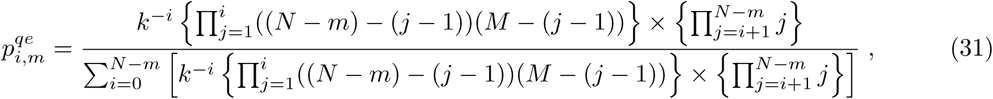

Finally, by defining

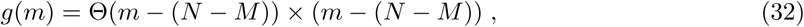

where Θ(*m* − (*N* − *M*)) is the Heaviside step function, we can write down a joint expression for all groups 0 ≤ *m* ≤ *N* and 0 ≤ *i* ≤ *M* − *g*(*m*):

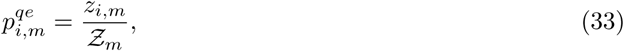

with

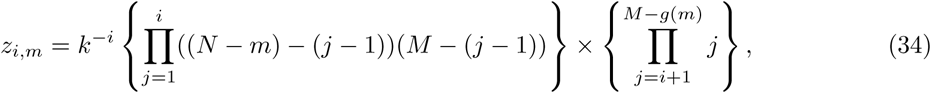

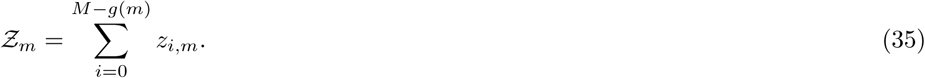

We now proceed to calculate 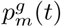. From Fig. 3, we observe that the transitions between the groups are described by the master equation identical in form to Eq. (10). However, the transition rates *a*_*m*_ in this case are different, as the group *m* can be reached from any of the *M* − *g*(*m* − 1) microstates in the group *m* − 1 (excluding only the microstate with *M* free enzymes) and we must also take into account the quasi-equilibrium probabilities of being in the corresponding microstate. It follows that the transition rates can be defined as:

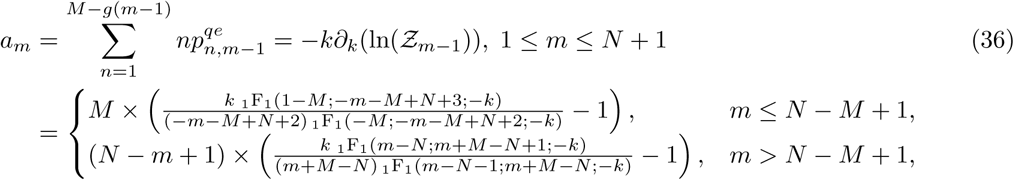

where _1_F_1_(*a, b, c*) is the confluent hypergeometric function. As the dynamics between the groups are identical to the single enzyme case, 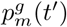 has exactly the same form as Eq. (21) but with the eigenvalues of 𝒬 being given by *λ*_*i*_ = − *a*_*i*_, where the *a*_*i*_ are now defined in Eq. (36).

We can now obtain the probability distribution *P* (*n*; *t*), which requires us to find all microstates in the system containing *n* free substrate molecules. From Fig. 3 we see that for substrate numbers *n*, where 0 ≤ *n* ≤ *N* − *M*, there are *M* + 1 corresponding microstates given by (*n*, 0), (*n*, 1), …, (*n, M*) which respectively belong to groups (*N* − *M*) −*n*, (*N* − *M*) −*n*+1, …, *N* − *n*. Therefore, the distribution has the form:

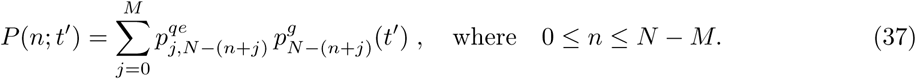

In the case of *N* − *M* < *n* ≤ *N*, there are *N* − (*n* − 1) microstates containing *n* substrate molecules, explicitly defined as (*n, M* − (*N* − *n*)), (*n, M* − (*N* − *n*) + 1), …, (*n, M*) and associated with groups 0, 1, …, *N* − *n* respectively. Hence we have:

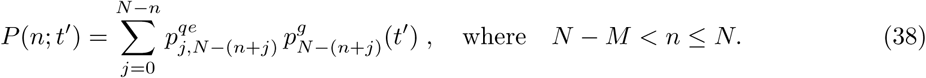

Finally, using the function *g*(*m*) previously defined in Eq. (32), we obtain:

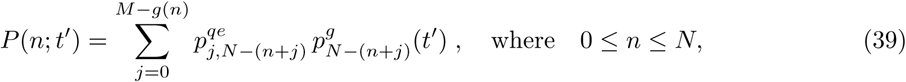

which fully describes the time-dependent solution for the multiple enzyme system *N* ≥ *M* with the initial condition 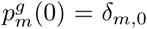. Note that the solution can also be extended to a more general initial distribution in the same way as was done for the single enzyme system in Section 3.1. The equations for mean number of substrate, ⟨*n*(*t*′)⟩, and standard deviation, σ(*t*′), at rescaled time *t*′ are the same as in Eqs. (23)-(24), but where *P* (*n*; *t*′) is now given by Eq. (39).

Now consider a multiple enzyme system which initially contains fewer free substrate molecules than enzymes, i.e., *M* ≥ *N*. The Markov chain describing the transitions between the microstates of this system, shown in Fig. 4, has similarities to that for the system with *N* ≥ *M*. Specifically, if we replace *N* by *M* in groups 0 to *N* in the *M* ≥ *N* case of Fig. 4 then we exactly recover groups *N* −*M* to *N* in the *N* ≥*M* case of Fig. 3. This mapping implies that the dynamics of the system with *M* ≥ *N* are correctly described by Eq. (39) due to the utility of *g*(*m*). Therefore, Eq. (39) is a valid solution for any positive integer values of *N* and *M*.

**Figure 4:**
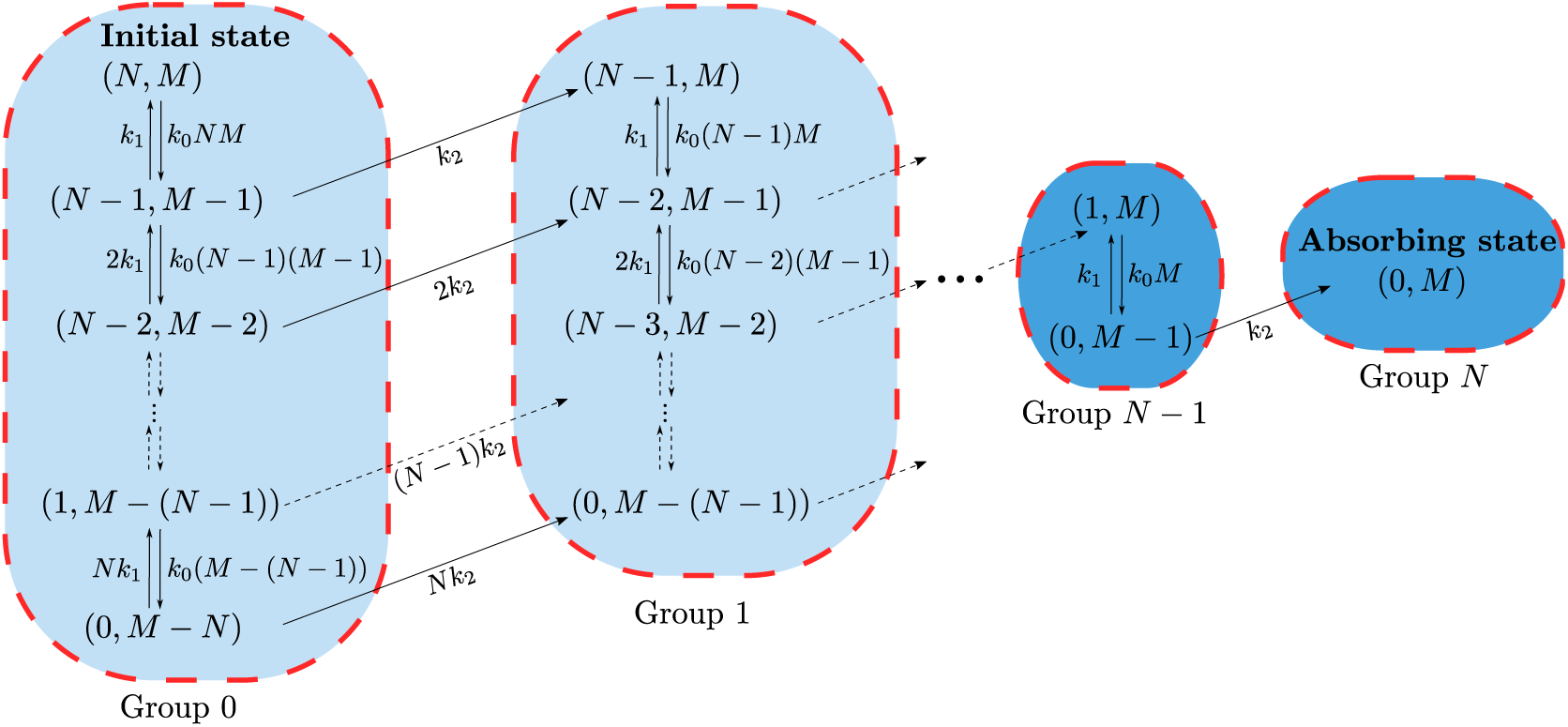
Illustration showing the transitions between the discrete microstates of the enzyme system (1) with initially *M* enzymes and *N* substrate molecules where *M* ≥ *N*. Fast processes are aggregated together, with each set of fast processes corresponding to a group. The dynamics of the groups 0 to *N* can be mapped onto the dynamics of groups *N* − *M* to *N* in the system with *N* ≥ *M* (shown in Fig. 3). See text for discussion.

As for the single enzyme case, we can make the initial statement that *k*_2_ must be sufficiently small for the derivation to hold, more precise. Suppose we are in the microstate (*n, n*_*e*_). There are then 3 possible reactions which can occur: (i) (*n, n*_*e*_) → (*n, n*_*e*_ + 1) with rate *k*_2_(*M − n*_*e*_), (ii) (*n*, − *n*_*e*_) → (*n* + 1, *n*_*e*_ + 1) with rate *k*_1_(*M* −*n*_*e*_) and (iii) (*n, n*_*e*_) → (*n* − 1, *n*_*e*_ − 1) with rate *k*_0_ *n n*_*e*_. Only the first reaction leads to a transition out of the current group of microstates (since its associated with the product formation step) and hence the probability of exiting the current group is *k*_2_(*M* −*n*_*e*_)*/*((*k*_1_ +*k*_2_)(*M* − *n*_*e*_)+*k*_0_ *n n*_*e*_). It is easy to prove that the latter is always less than *k*_2_*/*(*k*_1_+*k*_2_). Hence quasi-equilibrium of microstates in each group is possible when *k*_2_*/*(*k*_1_ + *k*_2_) ≪ 1. In other words, generally the closed-form solution for the distribution of substrate numbers will be accurate for all times provided *k*_1_≫ *k*_2_.

In Fig. 5A(i-iii) and 5B(i-iii) we show agreement between *P* (*n*; *t*′) from Eq. (39) and the SSA where *k*_1_ ≫ *k*_2_ is enforced, over times ranging between the initial time, when the number of substrate is *n* = *N* and the absorbing state at *n* = 0 for large times, for cases *M* ≥ *N* and *N* ≥ *M* respectively. In Fig. 5A(iv) and 5B(iv) we plot the mean and standard deviation of our analytical distribution (⟨*n*⟩, σ), the deterministic mean ⟨*n*⟩_*d*_ and the mean predicted by the SSA for *M ≥ N* and *N ≥ M* respectively. The SSA prediction of the mean is shown to be in exact correspondence with ⟨*n*⟩ when the QEA holds. Again, as previously discussed in Section 3.1, ⟨*n*⟩_*d*_ differs from ⟨*n*⟩ for small times due to the molecular discreteness of substrate. However, the discrepancy seen for ⟨*n*⟩ ⪅ 1 is no longer observed, highlighting that the discrepancy seen in Fig. 2A(iv) originates from the molecular discreteness of the enzyme species. We additionally note the presence of transient bimodality in Fig. 5B(ii) similar to that seen in the single enzyme case from Section 3.1; note that the parameter set chosen for Figs. 5A(i-iii) does not exhibit transient bimodality. The parameter space of transient bimodality is explored later in more detail in Section 3.2.2. In Fig. 6 we demonstrate using stochastic simulations that, as predicted by our theory, the requirement for the stochastic QEA to be a good approximation relies only on satisfying the condition *k*_1_ ≫ *k*_2_, and does not require any additional constraint on the value of *k*_0_.

**Figure 5:**
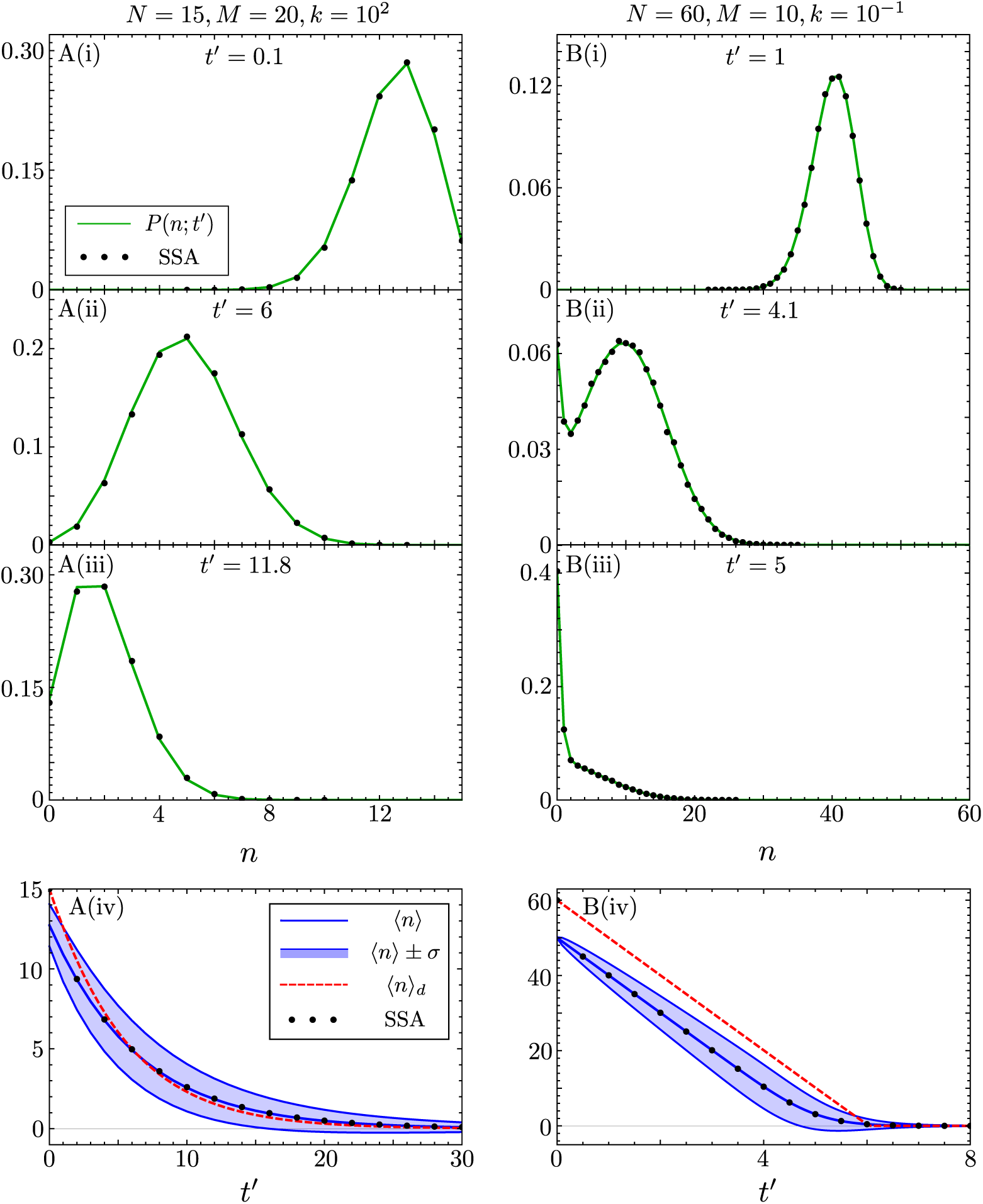
Comparison of the closed-form time-dependent probability distribution of substrate molecules, for the enzyme reaction (1) with multiple enzyme molecules *M*, and initial substrate molecules *N*, to the distribution obtained from the SSA. Note that the closed-form solution is given by Eq. (39). In A(i)-(iii), *N* = 15, *M* = 20, *k* = 10^2^ and we simulate the SSA using *k*_0_ = 1, *k*_1_ = 10^2^ and *k*_2_ = 1; the theory (green lines) agrees with the SSA since the quasi-equilibrium assumption is justified, i.e., *k*_1_*/k*_2_ ≫ 1. In B(i)-(iii), *N* = 60, *M* = 10, *k* = 10^−1^ and we simulate the SSA using *k*_0_ = 10^3^, *k*_1_ = 10^2^ and *k*_2_ = 1; again the theory is in agreement with the SSA since quasi-equilibrium is justified. Note that these results show that the theory accurately describes both the *N* ≥ *M* and the *M* ≥ *N* cases. In A(iv) and B(iv) we show the corresponding plots of the time-evolution of the mean ⟨*n*⟩ and of the standard deviation σ of the distributions of substrate molecules, as predicted by our theory; these are compared with the mean calculated from the SSA and from the deterministic rate equation ⟨*n*⟩_*d*_ given by Eq. (6). The parameter set in B is shown to be transiently bimodal in B(ii), whereas for the parameter set describing A transient bimodality is not observed. Each SSA probability distribution here is constructed from 10^5^ individual reaction trajectories.

**Figure 6:**
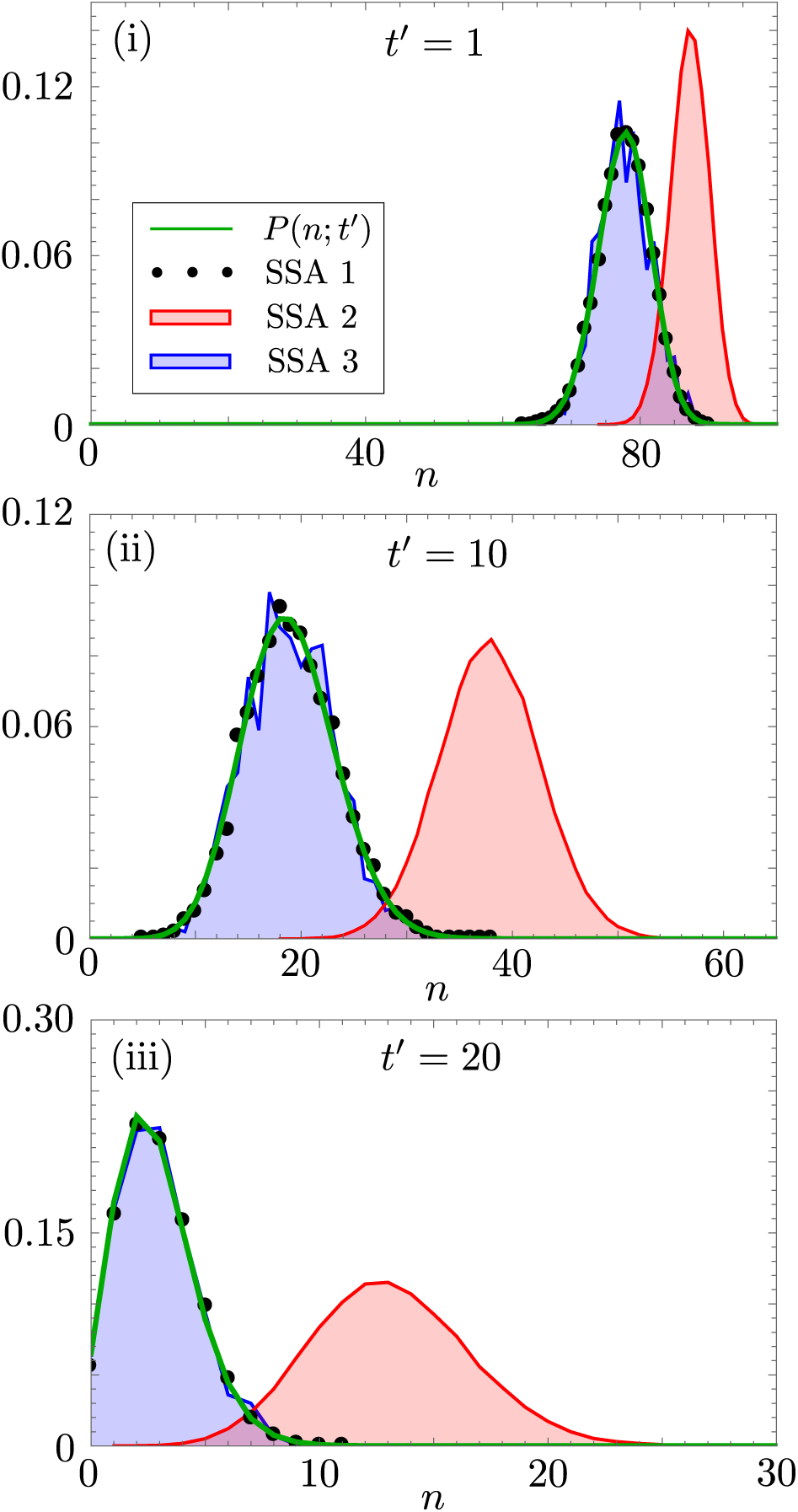
Testing the conditions necessary for the accuracy of the stochastic QEA. The three panels show how the accuracy of the closed-form time-dependent solution changes as we vary *k*_0_*/k*_2_ and *k*_1_*/k*_2_ whilst keeping *k* = *k*_1_*/k*_0_ fixed to 10^2^, for initial substrate number *N* = 10^2^ and total number of enzyme molecules equal to *M* = 25. The green line denotes the stochastic QEA solution from Eq. (39); SSA 1 (black dots) denotes the SSA prediction with parameters *k*_0_*/k*_2_ = 1, *k*_1_*/k*_2_ = 10^2^ calculated over 10^4^ trajectories; SSA 2 (blocked red region) denotes the SSA prediction with parameters *k*_0_*/k*_2_ = 10^−2^, *k*_1_*/k*_2_ = 1 calculated over 10^5^ trajectories; SSA 3 (blocked blue region) denotes the SSA prediction with parameters *k*_0_*/k*_2_ = 10, *k*_1_*/k*_2_ = 10^3^ calculated over 10^3^ trajectories. It is clear that SSA 2 is poorly predicted by *P* (*n*; *t*), which is expected as *k*_1_ = *𝒪*(*k*_2_). Since *P* (*n*; *t*) is in equally good agreement with SSA 1 and SSA 3 it can be seen that the only requirement is *k*_1_ ≫ *k*_2_, without requiring additional constraints on *k*_0_.

#### 3.2.1 Time-dependent solution for the probability distribution of enzyme molecules

Having solved the master equation for the group dynamics, it is relatively straightforward to extract the time-dependent probability distribution for the number of free enzyme molecules, *P* (*n*_*E*_; *t*′), and hence the distribution for the number of enzyme-substrate complexes, *P* (*n*_*C*_; *t*′). As previously, we begin by considering the *N* ≥ *M* system depicted in Fig. 3. We observe that the groups 0 ≤ *m* ≤ *N* − *M* all contain a microstate with *n*_*E*_ free enzyme molecules, where 0 ≤ *n*_*E*_ ≤ *M*, as enzymes are saturated with substrate. However, for groups *N* − *M* < *m* ≤ *N*, free enzymes become more abundant than free substrate molecules, so that microstates containing 0 < *n*_*E*_ ≤ *M* enzymes are found only in groups *N* − *M* < *m* ≤ *N* − (*M* − *n*_*E*_). Note that the quasi-equilibrium probability of having *n*_*E*_ free enzymes in group *m* is 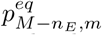, given by Eq. (33), and the group probabilities 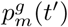 are identical to the ones defined for the distribution of substrate number in Eq. (39). Therefore, the distribution of free enzymes takes the form:

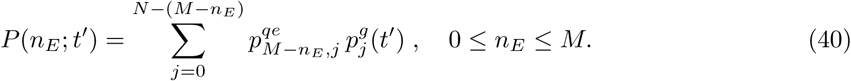

This expression is valid for any positive integer values of *N* and *M*, again due to the mapping between the Markov chains of *N* ≥*M* and *M* ≥*N* systems, described above. Moreover, for the *N* ≤*M* system, the definition of an empty sum as zero ensures that non-physical values of *n*_*E*_ are not allowed, i.e., the number of bound enzymes cannot be larger than *N* given the chosen initial conditions, so that *P* (*n*_*E*_; *t*′) = 0 for *n*_*E*_ < *N*. Finally, as *n*_*C*_ = *M n*_*E*_, the probability distribution of the enzyme-substrate complex follows trivially:

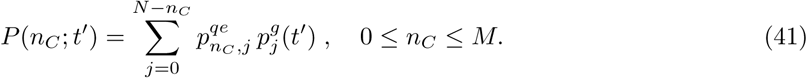

In Fig. 7A(i-iii) and 7B(i-iii) we confirm that *P* (*n*_*C*_; *t*′) from Eq. (41) and the SSA are in good agreement for enzyme systems with *M* ≥*N* and *N* ≥*M* respectively over the whole time-range from near the initial condition to the absorbing state, where again *k*_1_ ≫ *k*_2_ is enforced (using the same parameters as in Fig. 5). Note that the transient bimodality is seemingly not manifest in *P* (*n*_*C*_; *t*′) at the points in the parameter space where it is observed for the distribution of substrate number (c.f. Fig. 5B(ii) and 7B(ii)). In Fig. 7A(iv) and Fig. 7B(iv) we plot the mean and standard deviation of our analytical distribution for the enzyme-substrate complexes (⟨*n*_*C*_⟩ and *σ*_*C*_), and the mean predicted by the SSA for *M* ≥*N* and *N* ≥*M* respectively. The SSA prediction of the mean matches ⟨*n*_*C*_⟩ for all times further validating our solution, given that the QEA condition holds.

**Figure 7:**
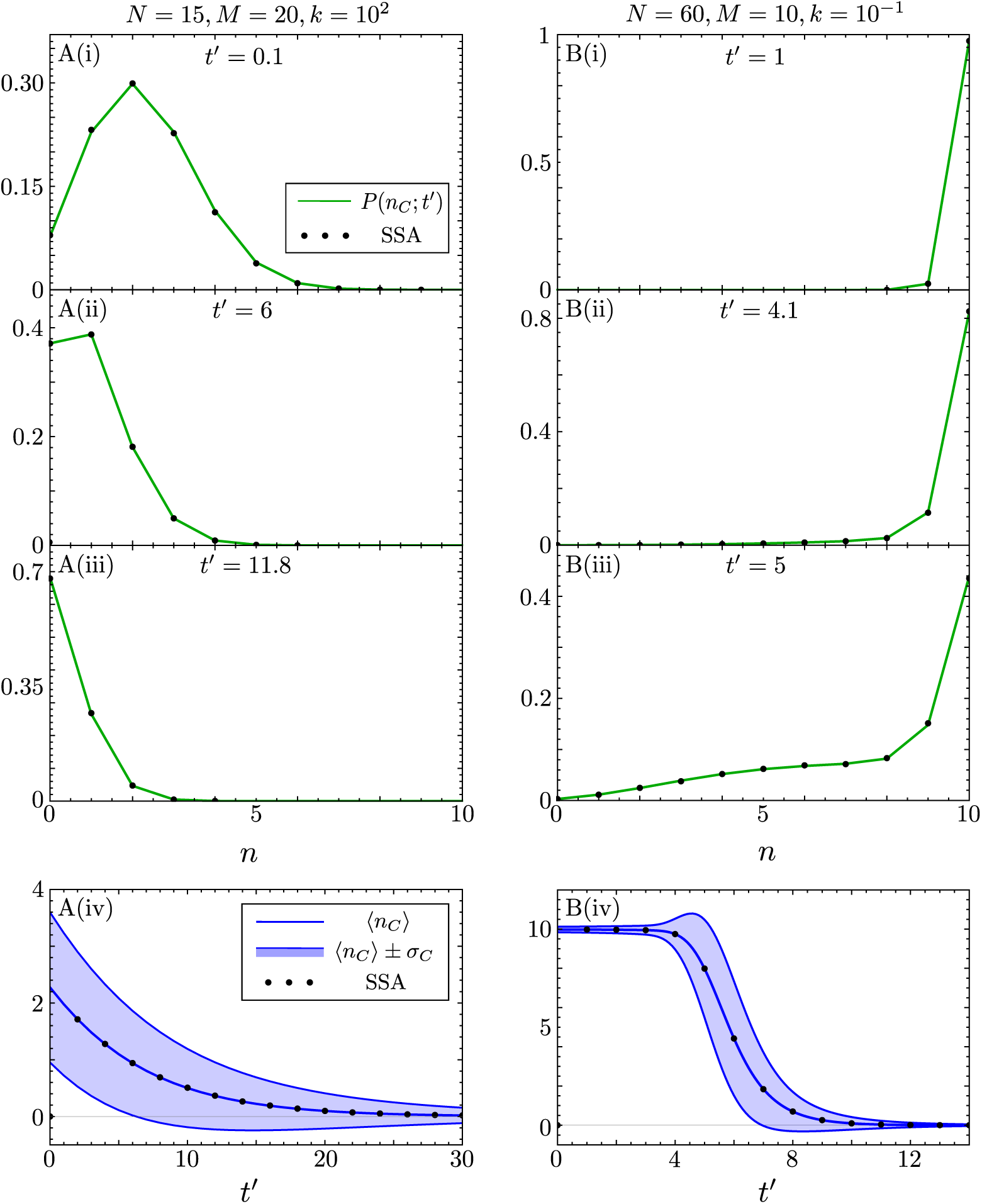
Comparison of the closed-form time-dependent probability distribution of enzyme-substrate complexes, for the enzyme reaction (1) with multiple enzyme molecules *M*, and initial substrate molecules *N*, to the distribution obtained from the SSA. Note that the closed-form solution is given by Eq. (41). In A(i)-(iii), *N* = 15, *M* = 20, *k* = 10^2^ and we simulate the SSA using *k*_0_ = 1, *k*_1_ = 10^2^ and *k*_2_ = 1; In B(i)-(iii), *N* = 60, *M* = 10, *k* = 10^−1^ and we simulate the SSA using *k*_0_ = 10^3^, *k*_1_ = 10^2^ and *k*_2_ = 1 (parameters are the same as in Fig. 5). In both cases, the theory (green lines) agrees with the SSA since the quasi-equilibrium assumption is justified, i.e., *k*_1_*/k*_2_ ≫ 1. In A(iv) and B(iv) we show the corresponding plots of the time-evolution of the mean ⟨*n*_*C*_⟩ and of the standard deviation σ_*C*_ of the distributions of enzyme-substrate complex, as predicted by our theory; these are compared with the mean calculated from the SSA. Each SSA probability distribution here is constructed from 10^5^ individual reaction trajectories.

#### 3.2.2 Bimodality

In Fig. 8A(i)-(iii) we explore further the transient bimodality observed in Figs. 2A(ii), 2B(iii) and 5B(ii). Namely, we investigate how the strength of the bimodality varies with the parameters *N, M* and *k* using the stochastic QEA solution from Eq. (39). Each point on the heatmap in Fig. 8A(i)-(iii) shows, for a particular parameter set, the maximum of the strength of bimodality calculated over the entire time course from *t*′ = 0 to a time near the absorbing state of *n* = 0. We utilise the measure of bimodality strength introduced in [39], which is explicitly given by:

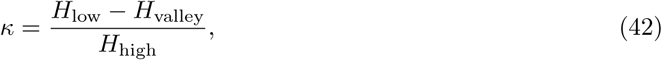

where *H*_low_ and *H*_high_ are the heights of the smallest and largest magnitude modes respectively, and *H*_valley_ is the height of the valley between the modes. For bimodal distributions *κ* has a value between 0 (no bimodality) and 1 (maximum bimodality), and for monomodal distributions is defined as zero. This definition of bimodality strength considers the ‘most bimodal’ distributions to have modes of equal height with a deep valley between them. In order to produce each heatmap we devised a simple algorithm, as follows. For each parameter set {*N, M, k*}:

**Figure 8:**
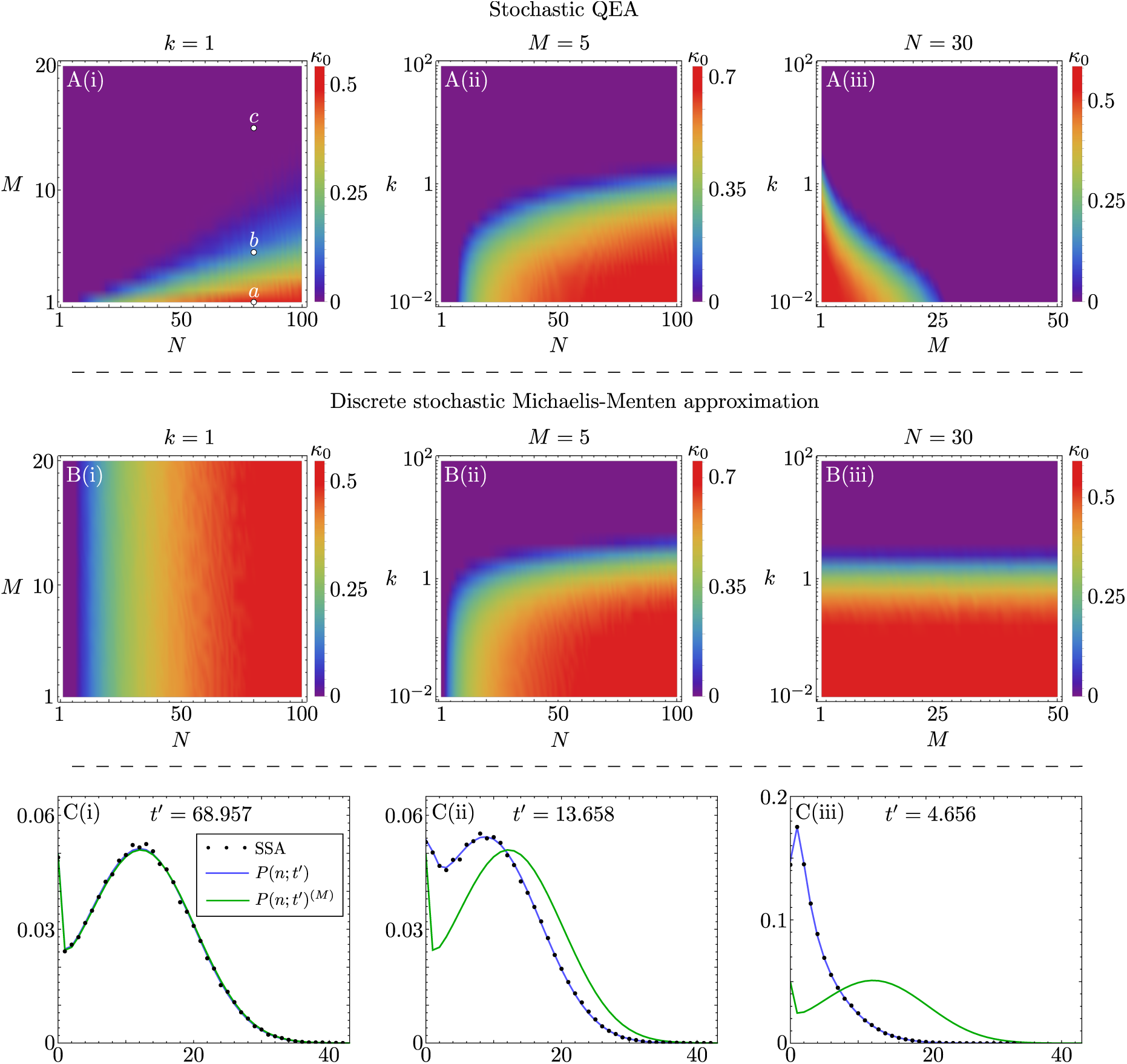
Heatmaps elucidating the regions of parameter space where transient bimodality is observed using the stochastic QEA solution (A(i)-(iii)) from Eq. (39) and the discrete stochastic MM approximation (B(i)-(iii)) given by Eq. (47). Note that *κ*_0_ is a measure of how bimodal is the distribution of substrate molecules across the timecourse of the reaction (see text for details). Three parameter regimes are considered: *N* vs *M* with *k* = 1 (left), *N* vs *k* with *M* = 5 (middle) and *M* vs *k* with *N* = 30 (right). The plots C(i)-(iii) show the closed-form distributions of the stochastic QEA, *P* (*n*; *t*′), and the discrete stochastic MM approximation, *P* (*n*; *t*′)^(*M*)^, at the times when the stochastic QEA exhibits maximum bimodality, for cases with *k* = 1, *N* = 80 and (i) *M* = 1, (ii) *M* = 5 and (iii) *M* = 15 (highlighted on the heatmap A(i) as the points *a, b* and *c* respectively). The corresponding SSA predictions with *k*_0_*/k*_2_ = 10^2^ and *k*_1_*/k*_2_ = 10^2^ are also included (constructed from 10^5^ individual reaction trajectories). Note that the two distributions (discrete stochastic MM approximation and stochastic QEA) are almost identical in C(i), but the difference becomes more pronounced in C(ii) and C(iii) with increasing *M*.

1. Calculate the estimated time to reach the absorbing state which provides us with the time range over which the transient bimodality search will be conducted. This can be estimated from the deterministic mean by solving Eq. (6) for *t*′ = *k*_2_*t* which gives us the required time range *T*_*a*_:

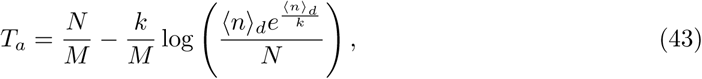

where we set ⟨*n*⟩_*d*_ = 10^−2^, which was chosen small enough such that transient bimodality for all parameter sets was accounted for.
2. Choose the number of iterations, *I*, over which to check if the distribution is bimodal. In our case we chose *I* = 400. This gives the set of times over which we check for bimodality as *t*_*i*_ = *iT*_*a*_*/I* for 1 ≤ *i* ≤ *I*.
3. Define a variable denoting the maximum bimodality measure *κ*_0_ which is initially set to zero. For each *t*_*i*_ find the number of peaks in the distribution given by Eq. (39) for the stochastic QEA, and if two peaks are detected, calculate the bimodality strength *κ* from Eq. (42). If *κ > κ*_0_ then set *κ*_0_ = *κ*. Do for all *t*_*n*_.
4. Once all iterations of this process are complete, the value of *κ*_0_ will denote the largest value of the transient bimodality measure for all probability distributions at *t* ∈ *t*_*i*_. We take *κ*_0_ as the largest value of transient bimodality encountered on the time course.

The results obtained using this algorithm are summarised by the three heatmaps in Fig. 8A(i)-(iii). The distribution of substrate molecules corresponding to the time at which the maximal bimodality strength *κ*_0_ occurs for points *a, b, c* in Fig. 8A(i) are shown by the solid blue lines in 8C(i)-(iii), respectively. Note that the bimodality is most pronounced in C(i), less in C(ii) and least in C(iii), in accordance with the value of *κ*_0_ in Fig. 8A(i); this validates the use of Eq. (42) as an accurate measure of the strength of bimodality. From Fig. 8A(i)-(iii), it is clear that bimodality is most pronounced when the initial number of substrate molecules *N* is significantly larger than the total enzyme number *M* and also when *k* is small, i.e., when the frequency of enzyme-substrate binding is much larger than the frequency of complex dissociation into enzyme and substrate. Note that generally the frequency of enzyme-substrate binding is inversely proportional to the volume of the compartment [6] in which the bimolecular reaction occurs and hence the transient bimodality is likely observable inside cells.

## 4 The discrete stochastic Michaelis-Menten approximation

We next consider how the analytical solution that we obtained for the reaction system (1) using a combination of averaging and linear algebra techniques in Section 3.2 compares with the solution of a commonly used reduced CME for enzyme kinetics.

The reduced CME for single substrate enzyme kinetics can be heuristically justified as follows (for a derivation see [12]). Under the QEA approximation, from the deterministic analysis in Section 2, it follows that the rate equation describing the time-evolution of the substrate concentration is given by:

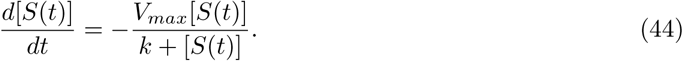

Note that *V*_*max*_ = *k*_2_*M*, where *M* is the total number of enzyme molecules. Hence, species *S* can be seen as changing into *P* by means of an effective first-order decay reaction with rate given by the right hand side of Eq. (44). One common way to approximately describe the enzyme reaction stochastically consists of writing down an effective propensity describing the decay of substrate, i.e., we postulate that if there are *m* substrate molecules at time *t* then the probability that a reaction *S* →*P* occurs somewhere in a unit volume in the time interval [*t, t* + *dt*) is approximately given by *a*_*m*_*dt* where *a*_*m*_ = *V*_*max*_*m/*(*k* + *m*). This is the discrete stochastic Michaelis-Menten (MM) approximation. Hence if we choose an initial condition of *N* substrate molecules, it follows that a corresponding effective CME is given by:

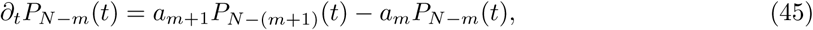

where *P*_*N*−*m*_(*t*) is the probability that there are *m* substrate molecules at time *t* (0 ≤*m* ≤*N*). This CME can be conveniently written as:

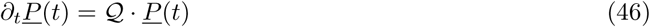

where *<u>P</u>* (*t*) = (*P*_0_(*t*), *P*_1_(*t*), …, *P*_*N*_ (*t*)) and 𝒬 is a (*N* + 1) × (*N* + 1) lower bidiagonal matrix whose only non-zero elements are 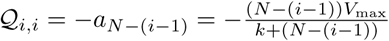 and 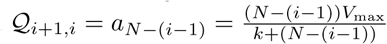 for 1 ≤ *i* ≤ *N* + 1. To our knowledge this master equation has not been analytically solved before, likely because a standard reference in the field of stochastic processes [7] states that for the one variable, one-step master equation, there are no general methods of solution except in the time-independent situation. However, using the method in [34] that was used to solve the master equation for the group dynamics for the single enzyme, the solution is found to be given by Eq. (21), modified to take into account the fact that *P*_*N*−*n*_ is equivalent to the probability of being in the group *N* − *n*:

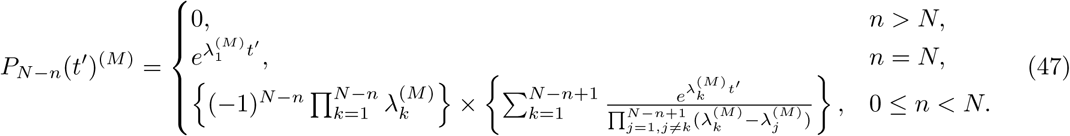

Note the superscript (*M*) specifying that the solution is for the CME (46) resulting from the discrete stochastic MM approximation. Here, we have again rescaled the time *t*′ = *k*_2_*t*, and 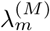 are the eigenvalues of 𝒬, which are simply given by the diagonal elements:

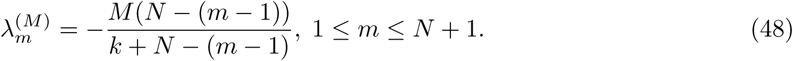

We shall denote the time-dependent mean and standard deviation of the distribution Eq. (47) by ⟨*n*(*t*′)⟩^(*M*)^ and σ(*t*′)^(*M*)^, respectively. Note that the distributions for the number of free enzymes/enzyme-substrate complexes cannot be obtained under the discrete stochastic MM approximation as the enzyme number fluctuations are not taken into account, in contrast to the Stochastic QEA from which enzyme/enzyme-substrate complex distributions can be obtained (see Section 3.2.1).

### 4.1 Comparison with the stochastic QEA

We used the algorithm described in Section 3.2.2 (with the difference that in step 3 we use Eq. (47) instead of Eq. (39)) to explore the regions of parameter space where the discrete stochastic MM approximation predicts the distribution of substrate molecules to be bimodal. The results are summarised by the three heatmaps in Fig. 8B(i)-(iii). By comparison to the heatmaps generated using the stochastic QEA in Fig. 8A(i)-(iii), it is clear that the discrete stochastic MM approximation tends to predict bimodality where in reality there is none. Notably, the bimodality predicted by the discrete stochastic MM approximation is independent of *M* (see Figs. 8B(i) and B(iii)) since *M* only acts to scale the eigenvalues representing the system’s relaxation timescales in Eq. (48); in contrast, the stochastic QEA predicts bimodality which is strongly dependent on *M* (see Figs. 8A(i) and A(iii)). These issues with the discrete stochastic MM approximation are also clearly discernible in 8C(i)-(iii), where we compare the distribution of substrate molecule numbers predicted by this approximation (green line) with that predicted by the SSA (dots) and the stochastic QEA (blue line).

A different way to contrast the discrete stochastic MM approximation and the stochastic QEA involves comparing the eigenvalues of the transition matrix. In the single enzyme case where *M* = 1, one observes that the eigenvalues predicted by Eq. (48) exactly match the eigenvalues predicted by averaging for the group dynamics in the single enzyme case from Eq. (11). However, note that the group dynamics is not precisely the same as the substrate dynamics which is determined by two microstates in different groups. For example the averaging technique implies that there are two microstates that contain *n* substrate molecules: (*n*, 0) and (*n*, 1) associated with groups *N* − (*n* + 1) and *N* − *n* respectively. However this subtlety is not important if *N* ≫ 1 and hence the CME resulting from the discrete stochastic MM approximation will practically lead to the same results as averaging for most cases of interest.

The comparison is more complicated in the case of multiple enzymes (*M >* 1) and abundant substrate *N* ≫ 1, which we explore in Fig. 9 (for *N* = 100 and *k* = 1), showing how the discrete stochastic MM approximate solution differs to that from averaging as the ratio *M/N* is increased. We first consider the case where *M/N* = 1*/*20, and we see that ⟨*n*⟩^(*M*)^ in Fig. 9A(i) is a good approximation of ⟨*n*⟩ for the time range of interest, i.e., from the initial state at *N* = 100 to a time *t*′ = 30 where both ⟨*n*⟩ ^(*M*)^ and ⟨*n*⟩ are small quantities. Note that the error in the standard deviation for this parameter set, shown in 9A(ii), is also small. The slight difference in the relaxation dynamics is corroborated by small differences in the eigenspectra of *λ*_*m*_ (given in Eq. (36) again noting that *λ*_*i*_ = −*a*_*i*_) and 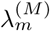 (given by Eq. (48)) which can be appreciated in Fig. (9)A(iii). We additionally plot the deterministic mean as predicted by Eq. (6) which clearly shows the relaxation dynamics of ⟨n⟩_*d*_ accurately approximates ⟨*n*⟩ for short times only.

**Figure 9:**
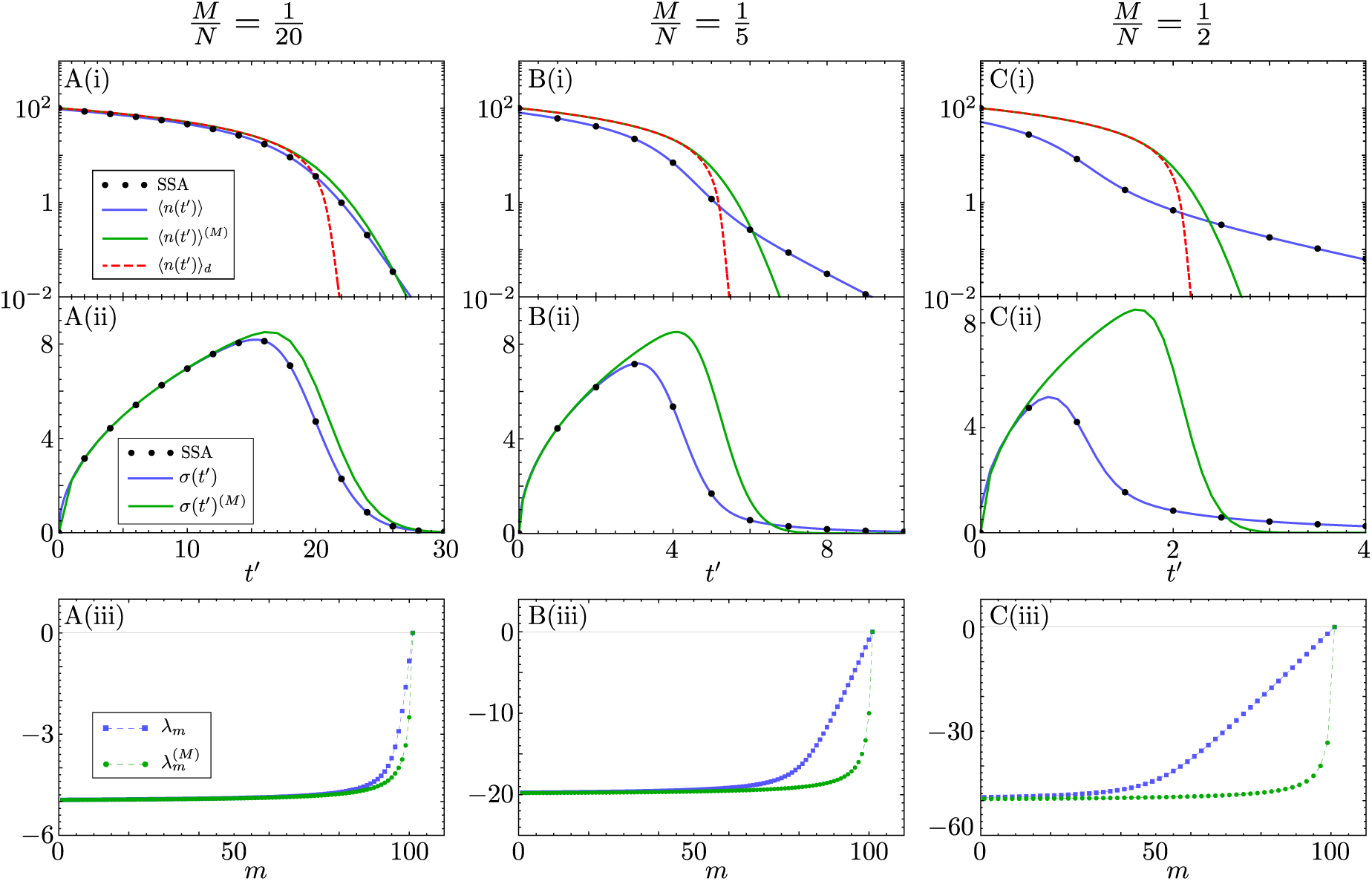
Comparison of the discrete stochastic MM approximation and the exact result from averaging in the quasiequilibrium limit. A(i), B(i) and C(i) show log-scale plots of ⟨*n*⟩, ⟨*n*⟩ ^(*M*)^ and ⟨*n*⟩ _*d*_ for *N* = 100, *k* = 1 and *M* = 5 (i.e, *M/N* = 1*/*20), *M* = 20 (i.e, *M/N* = 1*/*5) and *M* = 50 (i.e, *M/N* = 1*/*2) respectively. The corresponding SSA results with *k*_0_*/k*_2_ = 10^2^ and *k*_1_*/k*_2_ = 10^2^ are also included (constructed from 10^5^ individual reaction trajectories). A(ii), B(ii) and C(ii) are the corresponding plots of the standard deviations σ(*t*′), σ(*t*′)^(*M*)^ and that of SSA. A(iii), B(iii) and C(iii) show the eigenspectra for each differing *M/N*; each symbol corresponds to an individual eigenvalue (since the spectra are discrete) and the dashed lines are only present to aid the reader.

In Figs. 9B(i) and C(i) we see that as *M/N* increases to 1*/*5 and 1*/*2 respectively, ⟨*n*⟩^(*M*)^ becomes a worse approximation of ⟨*n*⟩, with ⟨*n*⟩^(*M*)^ tending more to ⟨*n*⟩_*d*_ than ⟨*n*⟩. The corresponding error in the standard deviation, as shown in 9B(ii) and C(ii), also follows that of the mean, increasing with *M/N*. There are two main reasons for this disagreement:

1. If *M* is comparable to *N* then initially there will be large fluctuations in the number of enzyme molecules, which are taken into account by the averaging solution (since it allows for switching between microstates in each group) but not by the CME resulting from the discrete stochastic MM approximation (since the total number of enzymes only appears as a constant through *V*_*max*_). This is most clearly seen in Fig. 9C(i) where we observe a large discrepancy between ⟨*n*⟩ and ⟨*n*⟩ (*M*) at 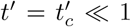 (where 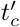 is the time over which the initial transient occurs and is indistinguishable from *t*′ = 0 in the figure).
2. Where *M/N* ≈ 𝒪(1), the eigenspectra *λ*_*m*_ and 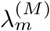 show a large disagreement (see Figs. 9B(iii) and C(iii)). This leads to the misprediction of the relaxation dynamics of ⟨*n*⟩^(*M*)^, which better represents the dynamics predicted by ⟨*n*⟩_*d*_ rather than of ⟨*n*⟩, for both small and large times. This is due to the fact that the effective Michaelis-Menten propensity in the reduced CME Eq. (46) is of the same form as the effective rate from the deterministic rate equation given by Eq. (44).

In summary, the solution of the CME obtained by the discrete stochastic MM approximation is a good approximation to the solution of the CME derived by averaging provided *N* ≫ 1 and *N/M* ≫ 1.

## 5 Multi-substrate mechanisms

Thus far we have considered the simple enzyme mechanism shown in (1) where an enzyme can catalyze a single type of substrate. However in nature, it is common for one enzyme species to be able to catalyze multiple substrates [47]. Multi-substrate reactions follow various mechanisms that describe how substrates bind and in what sequence. One such common mechanism is that of ternary complex formation, whereby two substrates bind sequentially to an enzyme to form a complex with three molecules. An example is the following mechanism involving two substrate species *A* and *B* and two corresponding reaction products, *P* and *Q* [47]:

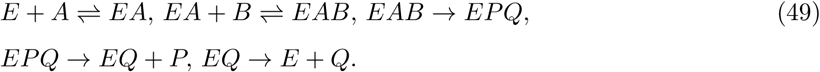

Note that here we have assumed an ordered binding mechanism, in the sense that binding of *A* must precede that of *B*. An alternative is a random binding mechanism, wherein either *A* or *B* could first bind the enzyme. We assume that both enzyme-substrate binding reactions and the steps subsequent to complex formation are fast such that we can consider the simpler reaction scheme:

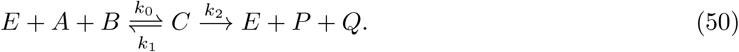

Note that ordered or random binding mechanisms cannot be distinguished within this reaction scheme. We assume that there are initially *N*_*A*_ molecules of substrate *A, N*_*B*_ molecules of substrate *B*, where *N*_*A*_ ≥*N*_*B*_, and *M* free enzymes. There exists a relation between the number of species *A* and *B*, denoted *n*_*A*_ and *n*_*B*_ respectively, which we can write as *n*_*A*_ −*n*_*B*_ = *N*_*A*_− *N*_*B*_ *≡* Δ_*AB*_. Hence each microstate of the system is fully specified by (*n*_*B*_, *n*_*E*_). Again the group dynamics where *k*_1_ ≫ *k*_2_ are given by Eq. (21) but the eigenvalues *λ*_*m*_ specific to this mechanism are given by:

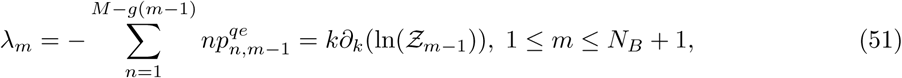

where we have now defined

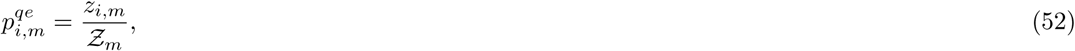

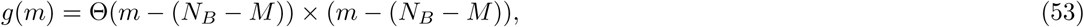

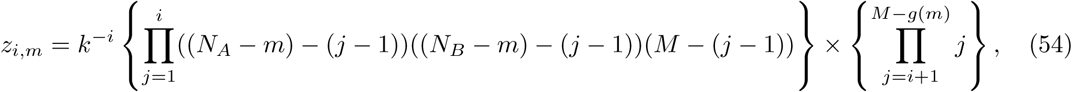

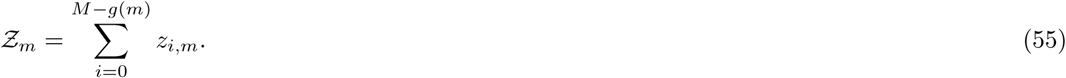

Using the results for the group dynamics and quasi-equilibrium probabilities, we can then find the probability distribution for the substrate molecules:

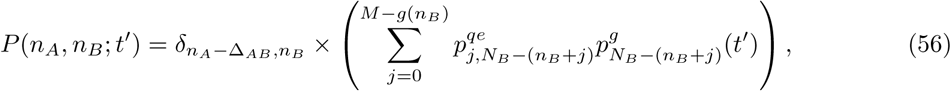

where *δ*_*i,j*_ is the Kronecker delta symbol. This allows us to find the marginal distributions:

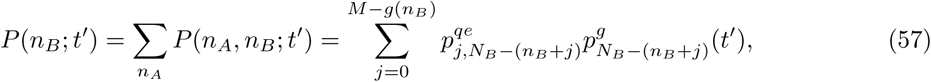

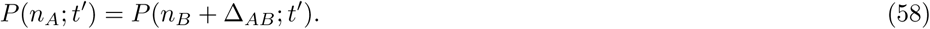

In Fig. 10 we compare the analytic marginal distributions against the SSA and as expected we find very good agreement when the rate parameters are consistent with the QEA. As previously for the single substrate mechanism, the distributions of *A* and *B* molecules display bimodality at intermediate times.

**Figure 10:**
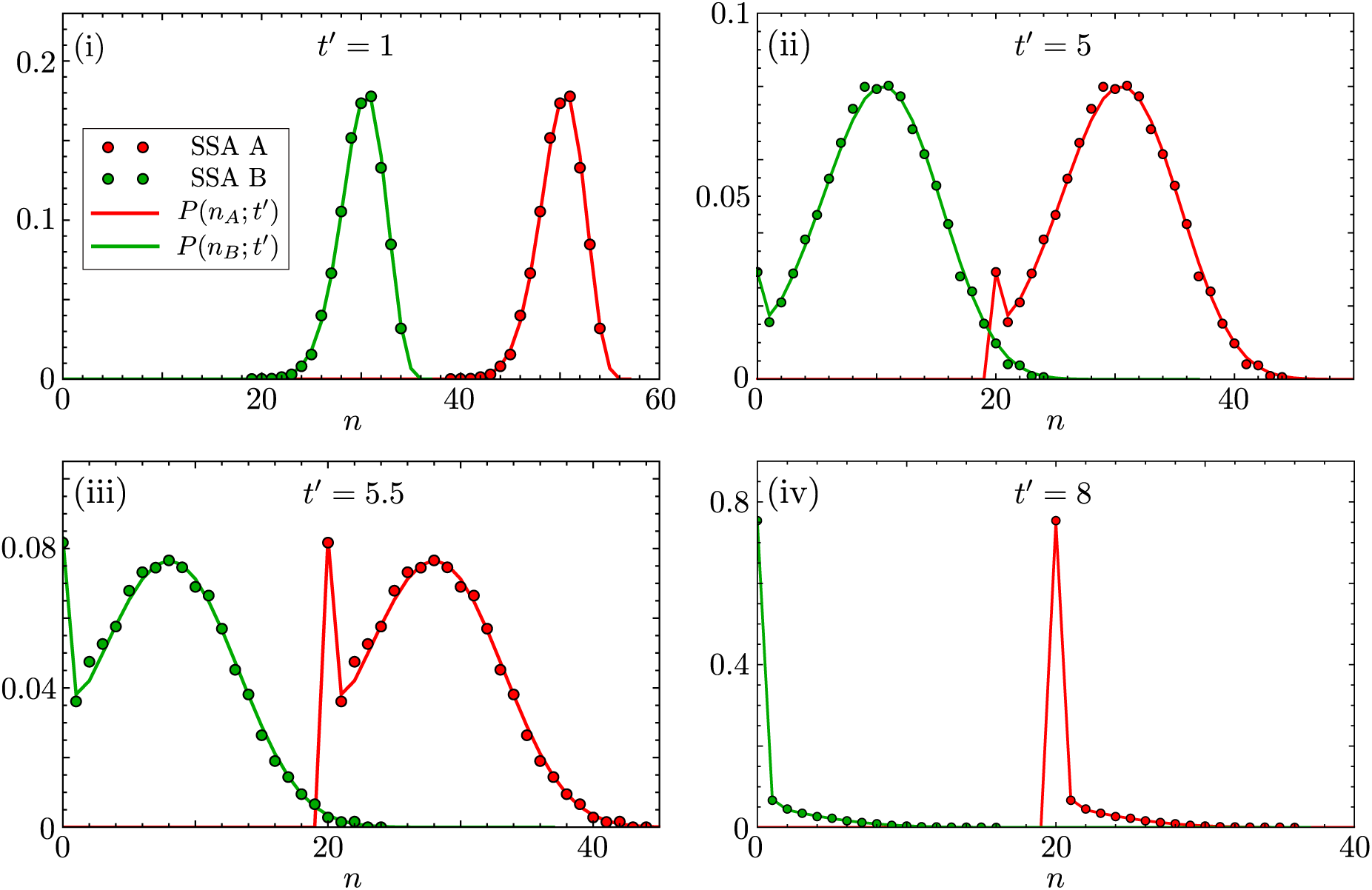
Comparison of the analytic distribution of two types of substrate species *A* and *B*, involved in the reaction mechanism (50), against the distributions obtained using the SSA. Note that SSA A and SSA B denote the SSA predictions for species A (red dots) and species B (green dots), respectively. We plot the probability distribution *P* (*n*_*A*_; *t*′) (red line; from Eq. (58)) and *P* (*n*_*B*_; *t*′) (green line; from Eq. (57)) for four different time points (time is non-dimensional as in previous figures). The initial number of substrate molecules are *N*_*A*_ = 60, *N*_*B*_ = 40 and the number of enzyme molecules is *M* = 5; the rates are *k*_0_*/k*_2_ = *k*_1_*/k*_2_ = 10^3^ which enforce the QEA. The analytic distributions are in good agreement with the respective SSA distributions. Note that the absorbing point of *A* is *n*_*A*_ = 20 while that of *B* is *n*_*B*_ = 0; this is dictated by the difference between the initial number of substrate molecules *N*_*A*_ − *N*_*B*_ = 20. Each SSA probability distribution is constructed from 10^5^ individual reaction trajectories.

## 6 Discussion

In summary, we have shown using averaging that in the limit of quasi-equilibrium between substrate and the enzyme, it is possible to reduce the two variable stochastic description of the MM reaction to that of an effective one variable master equation describing the slow transitions between groups of microstates. This master equation is subsequently solved exactly, using methods from linear algebra and complex analysis, to obtain closed-form solutions for the time-dependent marginal distributions of substrate and enzyme numbers. We have shown theoretically, and verified by means of stochastic simulations, that the solutions for the time-dependent marginal distributions are accurate for all times, provided the probability of complex decay into substrate and enzyme is much larger than the probability of complex decay into product and enzyme. To our knowledge, this is the first approximate closed-form solution for the MM reaction for an arbitrary initial number of substrate and enzyme molecules; previous work derived closed-form solutions for the case of a single enzyme molecule [9, 10] or else considered reactions with multiple enzyme molecules but focused on deriving expressions for the turnover rate [25, 30, 33]. We have also shown how the same procedure can be used to obtain the solution of more complex enzyme mechanisms such as those involving the catalysis of multiple types of substrate by the same enzyme species.

For the MM reaction, we have compared our closed-form solution with that obtained by the solution of the CME reduced by means of the widely used discrete stochastic MM approximation [12], where the propensity for substrate decay has a hyperbolic dependence on the number of substrate molecules. If the initial substrate number *N* is not much larger than the total enzyme number *M*, but the rate constants satisfy the inequality *k*_1_ ≫*k*_2_, then the enzyme numbers fluctuations can be large, even though the rapid equilibrium approximation is valid. In this case, we show that the distribution predicted by the CME reduced using the discrete stochastic MM approximation is significantly different than the one obtained from stochastic simulations, whereas the solution provided by our theory accurately matches the simulations.

Using the closed-form solution for the time-dependent marginal probability distribution for substrate number, we have found that unexpectedly for a delta function (unimodal) initial condition, the distribution of substrate numbers can be bimodal at intermediate times, if the initial number of substrate molecules is significantly larger than the total number of enzyme molecules and provided the rate of complex decay into substrate and enzyme is much less than the rate of substrate and enzyme binding. We note that the latter rate in the CME formulation is inversely proportional to the compartment volume (since the encounter rate of two molecules decreases with increasing volume [6]), and hence our results imply that in the limit of small volumes (taken at constant initial number of substrate and enzyme molecules), bimodality of the distribution of substrate molecules is observable. This result is of particular relevance to understanding enzyme dynamics inside cells where the volume is very small. Our system with the initial conditions used, can then be interpreted as modelling the enzyme-mediated decay of substrate molecules, following the production (via translation) of a short burst of substrate molecules *N* at time *t* = 0, provided there is not another burst of substrate expression before the substrate decays; these conditions are common for many cells where protein production occurs sporadically in bursts of short duration [48, 49]. We emphasize that the presence of transient bimodality in the MM reaction system is particularly interesting since it has no deterministic counterpart.

## Data Availability Statement

Data sharing is not applicable to this article since no new data were created or analyzed in this study.

## Acknowledgments

This work was supported by a BBSRC EASTBIO PhD studentship for J.H., a Turing scholarship for A.S. and a Leverhulme Trust grant (RPG-2018-423) for R.G.

## A Exact time-dependent solution of single enzyme system

The master equation for a single enzyme molecule (given by Eq. (7)) was first solved by Arányi and Tóth [9]. As the original paper is rather difficult to find, we present the solution here. The authors used marginal probability generating functions

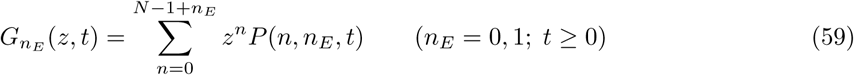

to transform Eq. (7) into the following first-order partial differential equations:

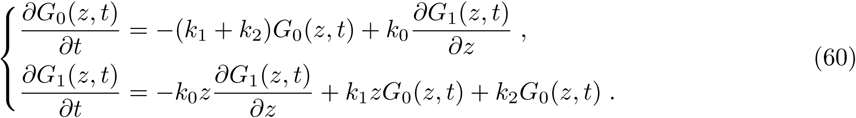

By a simple substitution one can prove that the solutions have the form:

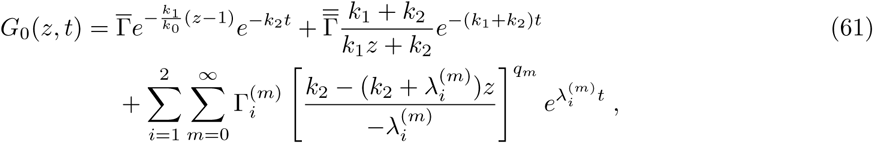

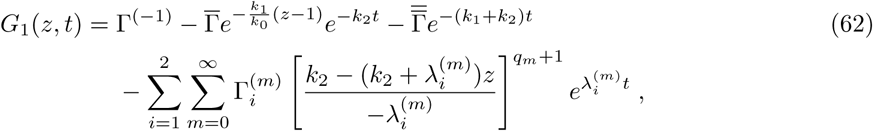

Where

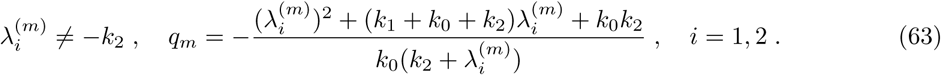

Since *G*_0_ and *G*_1_ are generating functions of a system with a finite state space, i.e., the number of substrate and enzyme are bounded quantities (*n* ∈ [0, *N*], *n*_*E*_ ∈ [0, 1]), they must be polynomials of a finite degree in *z*. Hence, <u>t</u>he summations in Eqs. (61) and (62) must contain a finite number of terms only, meaning that 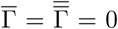 (if *k*_1_ ≠ 0). By the same reasoning the *q*_*m*_ must be positive integers, i.e., 0 ≤ *q*_*m*_ ≤ *N* − 1, (*q*_*m*_ = *m*), then the *λ*^(*m*)^ are the roots of a quadratic equation:

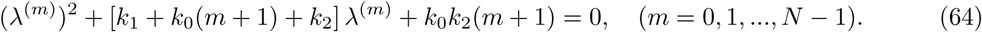

The constants Γ can be determined from the initial conditions:

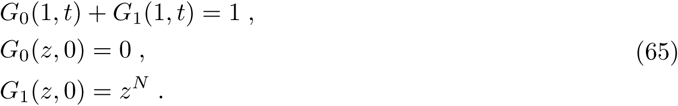

The first constraint implies that Γ^(−1)^ = 1, while the remaining two lead to a linear algebraic system for 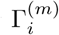 by enforcing the constraints explicitly on each coefficient of the polynomials *G*_0_ and *G*_1_ for each power of *z*. However, solving for 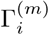 becomes computationally expensive for larger values of *N*.

To summarise, the solution has the form:

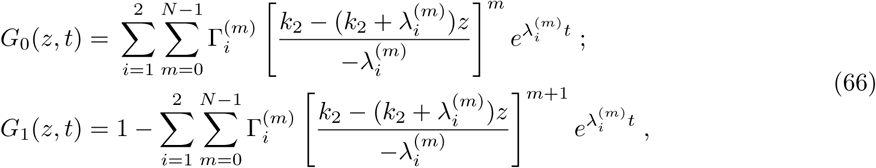

Where

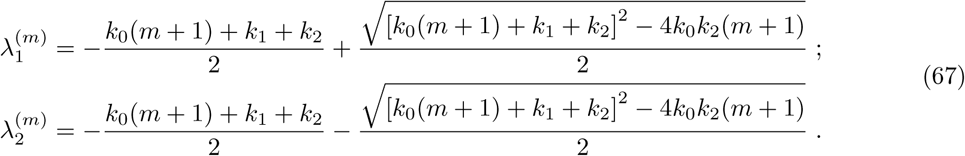

Finally, the probabilities can be calculated from the generating functions according to

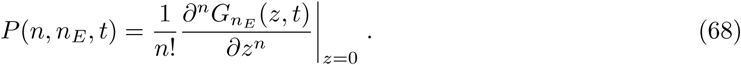

